# NETWORK-BASED FUNCTIONAL FRAGILITY REVEALS SYSTEM-LEVEL REORGANIZATION OF THE GUT MICROBIOME IN INFLAMMATORY BOWEL DISEASE

**DOI:** 10.64898/2026.04.16.719113

**Authors:** Mihika Kenavdekar, Elamathi Natarajan

**Affiliations:** Department of Bioinformatics, Biotecnika Info Labs Pvt Ltd, Bengaluru, India

**Keywords:** Gut microbiome, Inflammatory bowel disease, Functional networks analysis, Network topology, Keystone pathways, Machine learning, Microbiome reorganization

## Abstract

The human gut microbiome plays a critical role in host health, yet its functional organization in disease remains poorly understood. Most studies focus on taxonomic composition or pathway abundance, which fail to capture higher-order interactions governing system-level behavior.

Here, we investigated microbiome functional organization in inflammatory bowel disease (IBD), including Crohn’s disease (CD), ulcerative colitis (UC), and healthy controls (HC), using a network-based framework across 60 metagenomic samples. Functional pathway profiles were used to construct correlation-based interaction networks, followed by analysis of network topology, functional redundancy, keystone pathway architecture, and system robustness.

Disease-associated networks (CD and UC) exhibited reduced global connectivity, increased modular fragmentation, and centralization of keystone pathways, indicating a shift from distributed organization to more fragmented and fragile network structures compared to healthy controls. Notably, machine learning models demonstrated that network-derived features achieved higher classification performance (accuracy up to 0.824) compared to redundancy-based measures.

These findings reveal that microbiome dysfunction in IBD is driven by large-scale reorganization of functional interaction networks rather than loss of functional capacity. This study highlights the importance of network-level analysis in understanding microbiome-associated disease and provides a systems-level framework for future research.

## INTRODUCTION

The human gut microbiome constitutes a highly complex and dynamic ecosystem that plays a fundamental role in host physiology, metabolism, and immune regulation [1,2,3]. Disruptions in microbial communities have been strongly associated with a range of diseases, including IBD, which encompasses CD and UC [5,6,15,16]. Despite extensive investigation, the mechanistic basis of microbiome dysfunction in IBD remains incompletely understood [7,15]. Most existing studies have primarily focused on taxonomic composition or functional abundance to characterize microbial alterations in disease [11,12,16,29]. While these approaches have provided valuable insights, they inherently treat microbial functions as independent entities and fail to capture the interconnected nature of biological systems [8,27]. Emerging evidence suggests that microbial communities operate as coordinated networks, where interactions between functional pathways collectively determine system-level behavior [8,18,21]. Consequently, focusing solely on abundance may overlook critical structural changes that underlie disease-associated dysbiosis [9,21].

Recent advances in network biology offer a powerful framework to study such higher-order organization [27]. By modeling functional pathways as interconnected nodes within a network, it becomes possible to investigate how interactions, rather than mere presence or absence, are altered in disease states [8,19,20]. In this context, key properties such as network topology, centrality, modularity, and robustness provide insights into how microbial systems are organized and controlled [26,27]. However, the application of network-based approaches to microbiome functional data in IBD remains limited, and the extent to which disease is driven by structural reorganization versus functional loss is not fully resolved [9,21].

In parallel, the concept of functional redundancy has been proposed as a key determinant of microbiome stability, where multiple microbial pathways can compensate for one another to maintain system resilience [22,23,24]. Although redundancy is often assumed to buffer against perturbations, it remains unclear whether redundancy alone sufficiently explains microbiome dysfunction in complex diseases such as IBD or whether changes in interaction architecture play a more dominant role [22,24]. This limitation highlights a critical gap in current microbiome research, where functional presence is often conflated with functional organization [9,22].

In this study, we adopt an integrative systems-level approach to investigate microbiome functional organization across healthy controls and IBD conditions. Using metagenomic data from 60 samples (HC, CD, and UC), we constructed correlation-based functional interaction networks and systematically analyzed multiple dimensions of network behavior, including topology, centrality, modular structure, functional redundancy, keystone pathway organization, and robustness. Furthermore, we employed machine learning models to evaluate the relative predictive power of structural versus redundancy-based features in distinguishing disease states [13,14,28].

We hypothesize that microbiome dysfunction in IBD is not primarily driven by loss of functional components but rather by large-scale reorganization of interaction networks that govern system behavior [21,26]. By integrating network analysis, redundancy assessment, and predictive modeling, this study aims to provide a comprehensive understanding of how microbiome function is altered in disease, moving beyond abundance-based interpretations toward a structural and systems-oriented perspective.

## METHODOLOGY

### 3.1 DATA COLLECTION

A total of 60 publicly available metagenomic samples were obtained from the NCBI Sequence Read Archive (SRA) under BioProject accession PRJNA945504. The SRA is a widely used repository for high-throughput sequencing data [4]. The dataset included 30 HC, 15 CD, and 15 UC samples. Only fecal samples were selected to ensure consistency across all groups.

For each sample, corresponding BioSample identifiers, SRA run IDs, disease classification, and metadata such as sex were curated and organized into a structured dataset for downstream analysis.

**Table 1.**
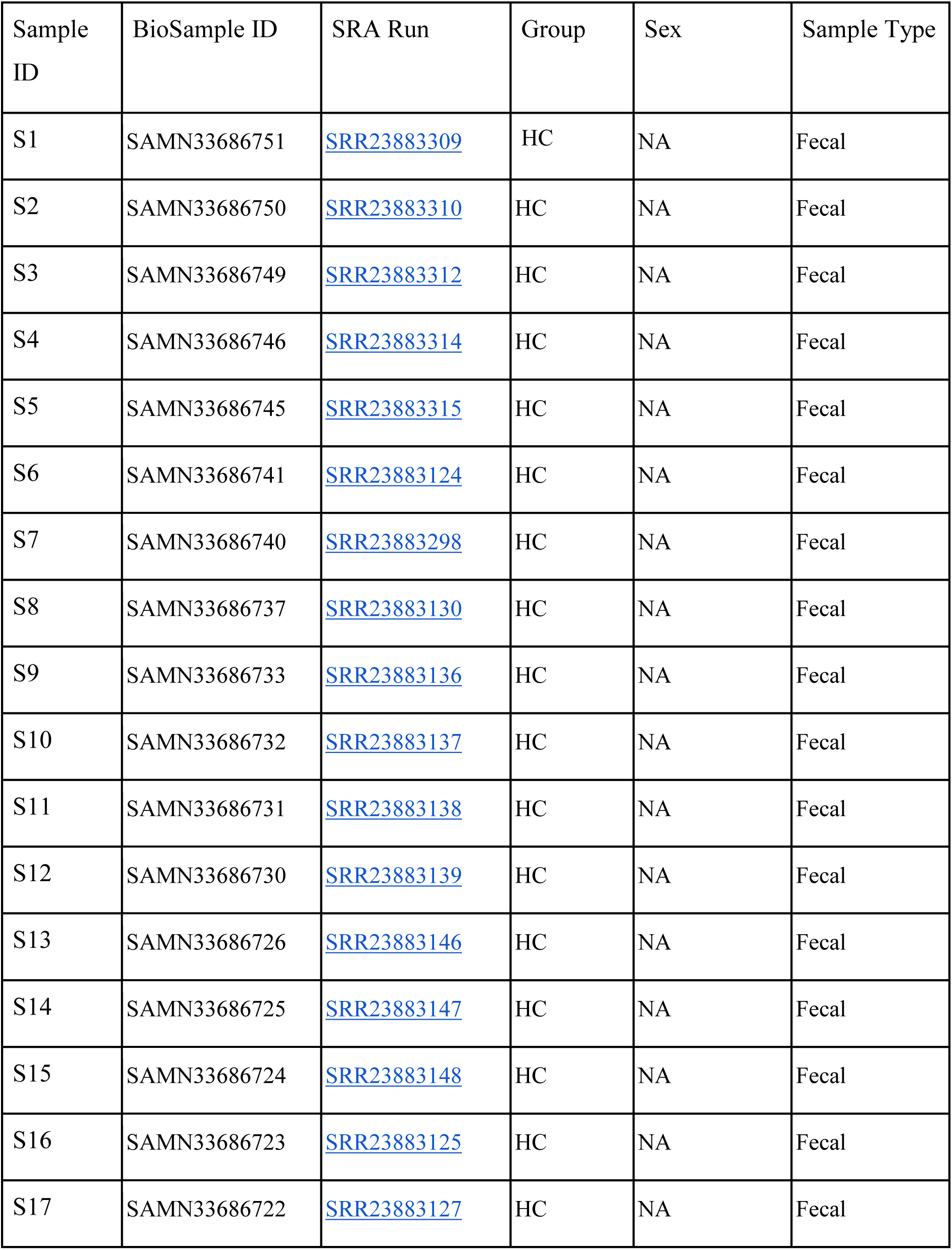
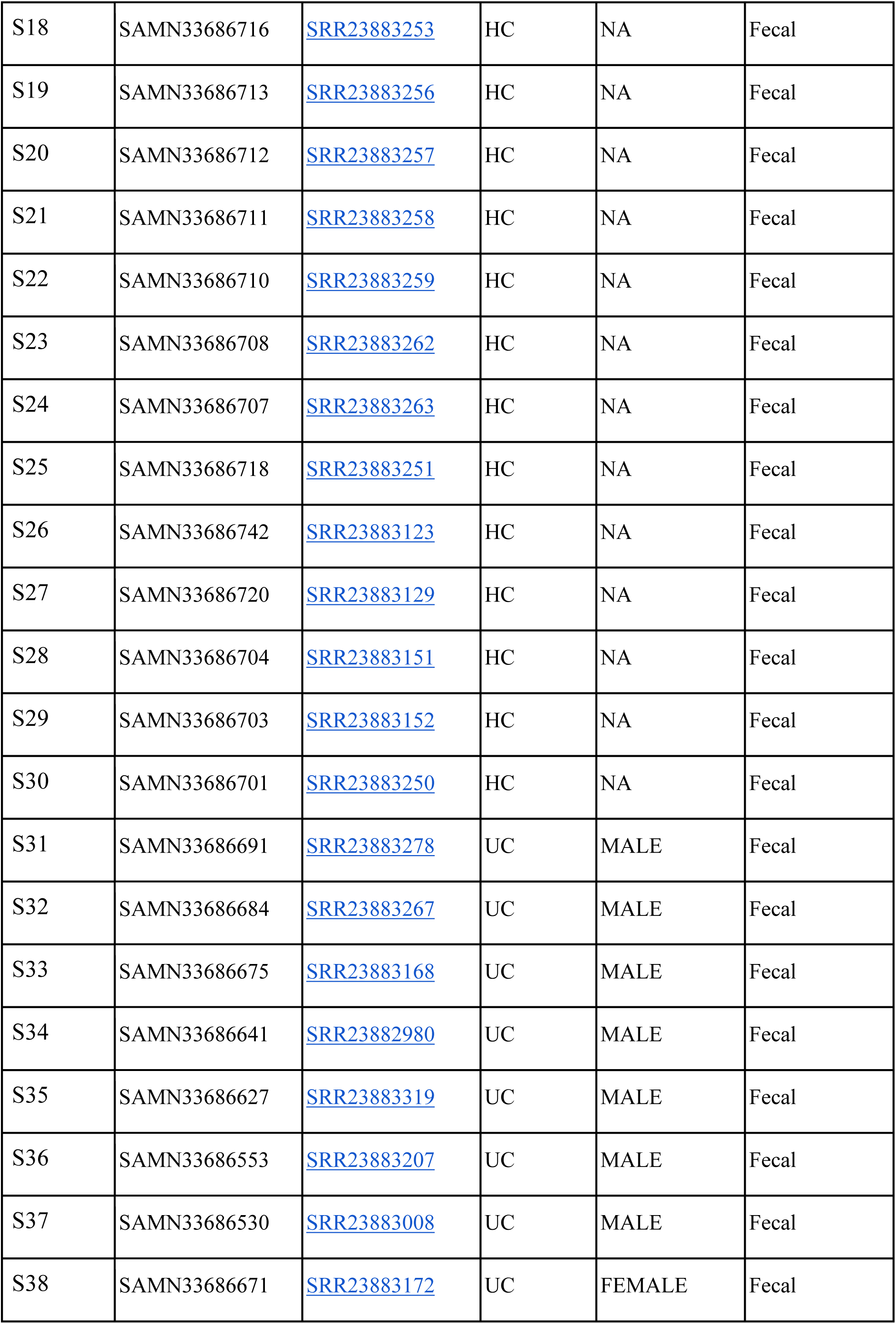
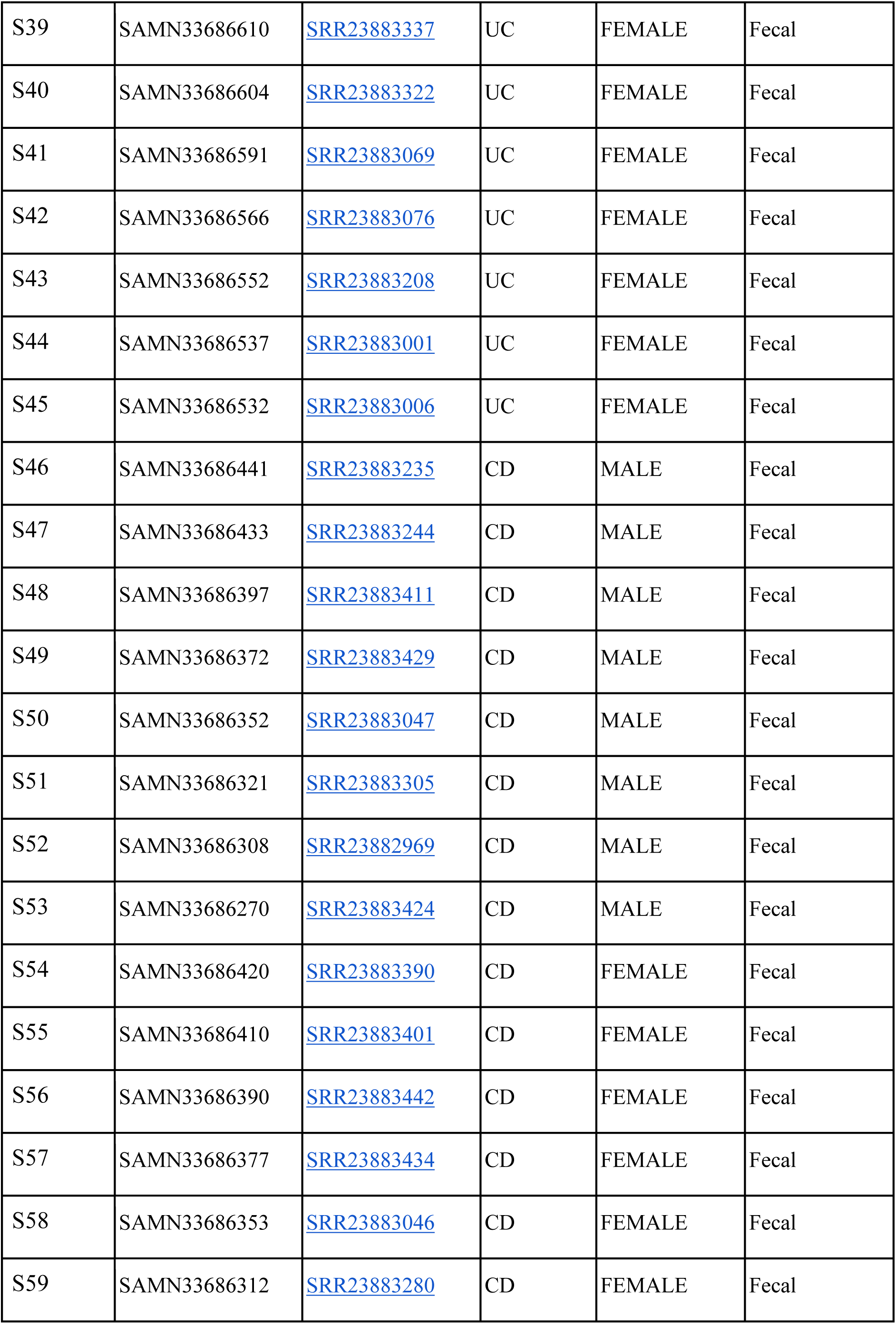
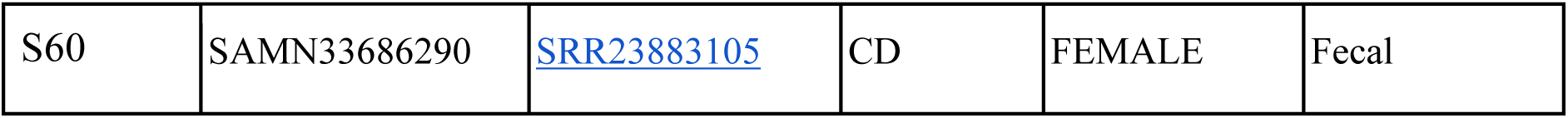
Summary of metagenomic samples used in this study.

| Sample ID | BioSample ID | SRA Run | Group | Sex | Sample Type |
| --- | --- | --- | --- | --- | --- |
| S1 | SAMN33686751 | SRR23883309 | HC | NA | Fecal |
| S2 | SAMN33686750 | SRR23883310 | HC | NA | Fecal |
| S3 | SAMN33686749 | SRR23883312 | HC | NA | Fecal |
| S4 | SAMN33686746 | SRR23883314 | HC | NA | Fecal |
| S5 | SAMN33686745 | SRR23883315 | HC | NA | Fecal |
| S6 | SAMN33686741 | SRR23883124 | HC | NA | Fecal |
| S7 | SAMN33686740 | SRR23883298 | HC | NA | Fecal |
| S8 | SAMN33686737 | SRR23883130 | HC | NA | Fecal |
| S9 | SAMN33686733 | SRR23883136 | HC | NA | Fecal |
| S10 | SAMN33686732 | SRR23883137 | HC | NA | Fecal |
| S11 | SAMN33686731 | SRR23883138 | HC | NA | Fecal |
| S12 | SAMN33686730 | SRR23883139 | HC | NA | Fecal |
| S13 | SAMN33686726 | SRR23883146 | HC | NA | Fecal |
| S14 | SAMN33686725 | SRR23883147 | HC | NA | Fecal |
| S15 | SAMN33686724 | SRR23883148 | HC | NA | Fecal |
| S16 | SAMN33686723 | SRR23883125 | HC | NA | Fecal |
| S17 | SAMN33686722 | SRR23883127 | HC | NA | Fecal |
| S18 | SAMN33686716 | SRR23883253 | HC | NA | Fecal |
| S19 | SAMN33686713 | SRR23883256 | HC | NA | Fecal |
| S20 | SAMN33686712 | SRR23883257 | HC | NA | Fecal |
| S21 | SAMN33686711 | SRR23883258 | HC | NA | Fecal |
| S22 | SAMN33686710 | SRR23883259 | HC | NA | Fecal |
| S23 | SAMN33686708 | SRR23883262 | HC | NA | Fecal |
| S24 | SAMN33686707 | SRR23883263 | HC | NA | Fecal |
| S25 | SAMN33686718 | SRR23883251 | HC | NA | Fecal |
| S26 | SAMN33686742 | SRR23883123 | HC | NA | Fecal |
| S27 | SAMN33686720 | SRR23883129 | HC | NA | Fecal |
| S28 | SAMN33686704 | SRR23883151 | HC | NA | Fecal |
| S29 | SAMN33686703 | SRR23883152 | HC | NA | Fecal |
| S30 | SAMN33686701 | SRR23883250 | HC | NA | Fecal |
| S31 | SAMN33686691 | SRR23883278 | UC | MALE | Fecal |
| S32 | SAMN33686684 | SRR23883267 | UC | MALE | Fecal |
| S33 | SAMN33686675 | SRR23883168 | UC | MALE | Fecal |
| S34 | SAMN33686641 | SRR23882980 | UC | MALE | Fecal |
| S35 | SAMN33686627 | SRR23883319 | UC | MALE | Fecal |
| S36 | SAMN33686553 | SRR23883207 | UC | MALE | Fecal |
| S37 | SAMN33686530 | SRR23883008 | UC | MALE | Fecal |
| S38 | SAMN33686671 | SRR23883172 | UC | FEMALE | Fecal |
| S39 | SAMN33686610 | SRR23883337 | UC | FEMALE | Fecal |
| S40 | SAMN33686604 | SRR23883322 | UC | FEMALE | Fecal |
| S41 | SAMN33686591 | SRR23883069 | UC | FEMALE | Fecal |
| S42 | SAMN33686566 | SRR23883076 | UC | FEMALE | Fecal |
| S43 | SAMN33686552 | SRR23883208 | UC | FEMALE | Fecal |
| S44 | SAMN33686537 | SRR23883001 | UC | FEMALE | Fecal |
| S45 | SAMN33686532 | SRR23883006 | UC | FEMALE | Fecal |
| S46 | SAMN33686441 | SRR23883235 | CD | MALE | Fecal |
| S47 | SAMN33686433 | SRR23883244 | CD | MALE | Fecal |
| S48 | SAMN33686397 | SRR23883411 | CD | MALE | Fecal |
| S49 | SAMN33686372 | SRR23883429 | CD | MALE | Fecal |
| S50 | SAMN33686352 | SRR23883047 | CD | MALE | Fecal |
| S51 | SAMN33686321 | SRR23883305 | CD | MALE | Fecal |
| S52 | SAMN33686308 | SRR23882969 | CD | MALE | Fecal |
| S53 | SAMN33686270 | SRR23883424 | CD | MALE | Fecal |
| S54 | SAMN33686420 | SRR23883390 | CD | FEMALE | Fecal |
| S55 | SAMN33686410 | SRR23883401 | CD | FEMALE | Fecal |
| S56 | SAMN33686390 | SRR23883442 | CD | FEMALE | Fecal |
| S57 | SAMN33686377 | SRR23883434 | CD | FEMALE | Fecal |
| S58 | SAMN33686353 | SRR23883046 | CD | FEMALE | Fecal |
| S59 | SAMN33686312 | SRR23883280 | CD | FEMALE | Fecal |
| S60 | SAMN33686290 | SRR23883105 | CD | FEMALE | Fecal |

### 3.2 DATA PREPROCESSING AND QUALITY CONTROL

Raw metagenomic sequencing data were downloaded from the NCBI Sequence Read Archive (SRA) using the SRA Toolkit (fasterq-dump v3.1.1) on the Galaxy Europe platform [4]. Quality assessment of raw sequencing reads was performed using FastQC (v0.11.9) and Falco (v1.2.4), a high-performance implementation of FastQC, to evaluate sequencing quality metrics including per-base quality scores, GC content, and adapter contamination.

Low-quality reads and adapter sequences were removed using fastp (v0.23.2) with default parameters to ensure high-quality input for downstream analysis. To eliminate host contamination, reads were aligned against the human reference genome (GRCh38) using Bowtie2 (v2.5.5), and mapped reads were discarded. This step ensured that only microbial sequences were retained for further analysis [20].

### 3.3 TAXONOMIC AND FUNCTIONAL PROFILING

Taxonomic profiling was performed using MetaPhlAn (v4.2.4), which identifies microbial taxa based on clade-specific marker genes [11]. Functional profiling was conducted using HUMAnN (v3.9), which maps microbial reads to known metabolic pathways and reconstructs pathway abundance profiles for each sample [11,12].

The resulting pathway abundance and coverage tables were used for downstream functional analysis. All profiles were normalized using relative abundance scaling to account for differences in sequencing depth across samples [12].

### 3.4 FUNCTIONAL NETWORK CONSTRUCTION

Functional interaction networks were constructed separately for each group (HC, CD, UC) using correlation-based approaches [18,19,20].

Pairwise associations between functional pathways were computed using Spearman rank correlation, chosen for its robustness to non-normal distributions commonly observed in microbiome data.

To define statistically significant interactions, correlations were filtered using the following criteria: absolute correlation coefficient threshold: |ρ| > 0.6 statistical significance: p < 0.05

To control for multiple hypothesis testing, Benjamini–Hochberg false discovery rate (FDR) correction was applied, and only associations with adjusted p < 0.05 were retained [19,20].

In these networks, nodes represent functional pathways, and edges represent statistically significant associations between pathways. This framework enables the investigation of higher-order functional organization within microbial communities [8,21].

Negative correlations were retained to capture both cooperative and competitive interactions.

### 3.5 NETWORK TOPOLOGY ANALYSIS

Network topology was characterized using standard graph-theoretical metrics, including degree centrality, betweenness centrality, and closeness centrality [27].

These measures were used to quantify node connectivity, control over information flow, and relative importance within the network [27].

All network metrics were computed using established graph analysis libraries in R (e.g., igraph package). Comparative analysis across HC, CD, and UC networks was performed to identify structural differences associated with disease states.

### 3.6 FUNCTIONAL REDUNDANCY ANALYSIS

Functional redundancy was quantified using pathway-level redundancy scores, representing the extent to which multiple pathways contribute to similar functional roles within the microbiome [22,23]. Redundancy distributions were compared across groups using non-parametric statistical tests (Wilcoxon rank-sum test) to assess differences in functional buffering capacity [24].

This analysis aimed to evaluate whether disease states are associated with reduced redundancy and diminished system resilience.

### 3.7 KEYSTONE PATHWAY IDENTIFICATION

Keystone pathways were identified based on centrality measures, including degree and betweenness centrality [10,25]. Pathways with centrality values in the top 10% of the distribution were classified as keystone candidates, representing critical regulators of network structure and function [25].

Comparative analysis across groups was performed to assess shifts in keystone pathway distribution and their role in disease-associated network reorganization.

### 3.8 ROBUSTNESS AND STABILITY ANALYSIS

Network robustness was evaluated by assessing the impact of node removal on overall network connectivity [26]. Simulated perturbations were performed using random node removal and targeted removal (based on high centrality nodes).

Network integrity was quantified using changes in the size of the largest connected component and network fragmentation metrics [26]. This analysis provided insights into the resilience of microbial functional networks under perturbation.

### 3.9 CORRELATION AND INTEGRATION ANALYSIS

The relationship between functional redundancy and network centrality was investigated using Spearman correlation analysis. Associations between redundancy scores and metrics such as degree and betweenness centrality were examined to understand how redundancy relates to network structure [22].

Statistical significance was assessed using p < 0.05, with FDR correction applied where appropriate.

### 3.10 MACHINE LEARNING ANALYSIS

Machine learning models were implemented to classify samples into CD, UC, and HC groups based on extracted functional features [14,28].

Three classification algorithms were employed: logistic regression, random forest, and support vector machines (SVM), enabling comparative evaluation across linear and nonlinear modeling approaches [14].

Two feature sets were evaluated:

i. network-derived features and
ii. redundancy-based features.

Feature scaling and normalization were applied where required prior to model training. Model performance was assessed using classification accuracy, confusion matrices, and receiver operating characteristic (ROC) curves. To ensure robustness and minimize overfitting, models were evaluated using 5-fold cross-validation [14].

Feature importance for the random forest model was assessed using mean decrease in accuracy and Gini index. This multi-model framework was used to evaluate the robustness, consistency, and predictive relevance of functional network organization in distinguishing disease states [28].

## RESULTS

To systematically investigate functional organization of the gut microbiome in health and disease, we performed a multi-level analysis integrating pathway abundance, network topology, functional redundancy, and predictive modeling. Results are presented sequentially from basic functional properties to higher-order structural and integrative analyses.

### 4.1 FUNCTIONAL REDUNDANCY ANALYSIS

To evaluate whether microbiome functional capacity is altered in disease, we first assessed functional redundancy across microbial pathways in healthy and IBD conditions. Functional redundancy reflects the extent to which multiple pathways contribute to similar biological functions, thereby indicating system resilience.

Comparison of redundancy distributions across groups revealed modest but statistically significant differences between HC and disease states (Wilcoxon test, p < 0.05). Healthy microbiomes exhibited relatively higher redundancy levels across several pathways, suggesting a more buffered functional system. In contrast, both CD and UC samples showed reduced redundancy in specific pathways.

However, the overall separation between groups remained limited, with substantial overlap observed across conditions (Figure 1). This indicates that although redundancy is affected in disease, its discriminatory power is weak and insufficient to fully explain microbiome dysfunction.

**Figure 1.**
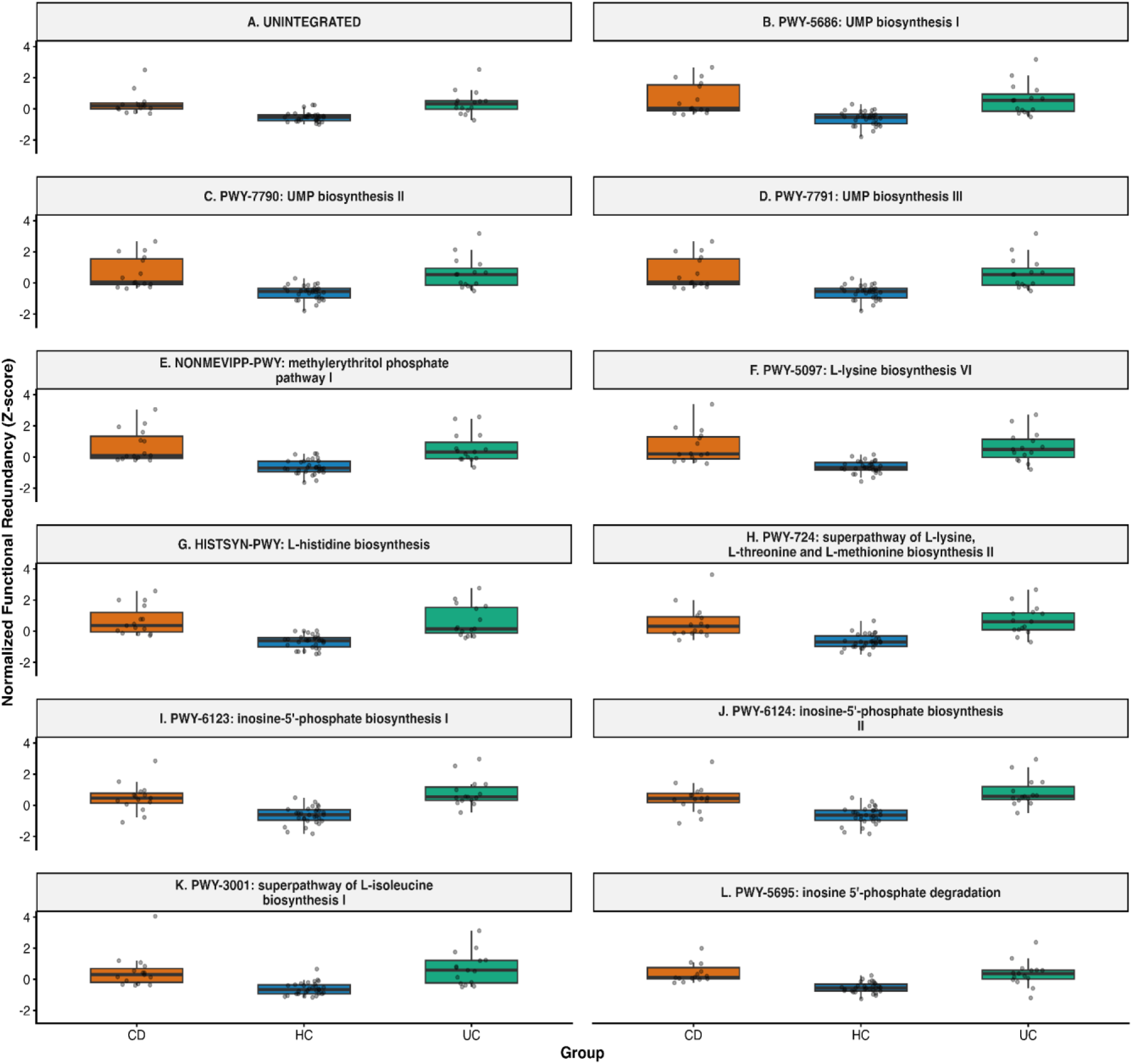
Functional redundancy across microbiome pathways in HC, CD, and UC groups. Boxplots represent the distribution of pathway-level redundancy scores across samples. While disease conditions show a trend toward reduced redundancy, overall group separation remains limited.

These findings indicate that while functional redundancy exhibits modest disease-associated changes, it does not capture the dominant mechanisms underlying microbiome dysfunction, highlighting the need to investigate higher-order interaction structures.

### 4.2 FUNCTIONAL COMPOSITION AND PATHWAY VARIATION

To investigate pathway-level functional variation across disease states, we analyzed the distribution of pathway abundances using hierarchical clustering and heatmap visualization. This approach enables identification of global functional patterns and potential disease-associated shifts in pathway composition.

Heatmap analysis revealed subtle but non-distinct differences in functional composition across groups. Hierarchical clustering did not result in clear separation of CD, UC, and HC samples, with substantial intermixing observed (Figure 2).

**Figure 2.**
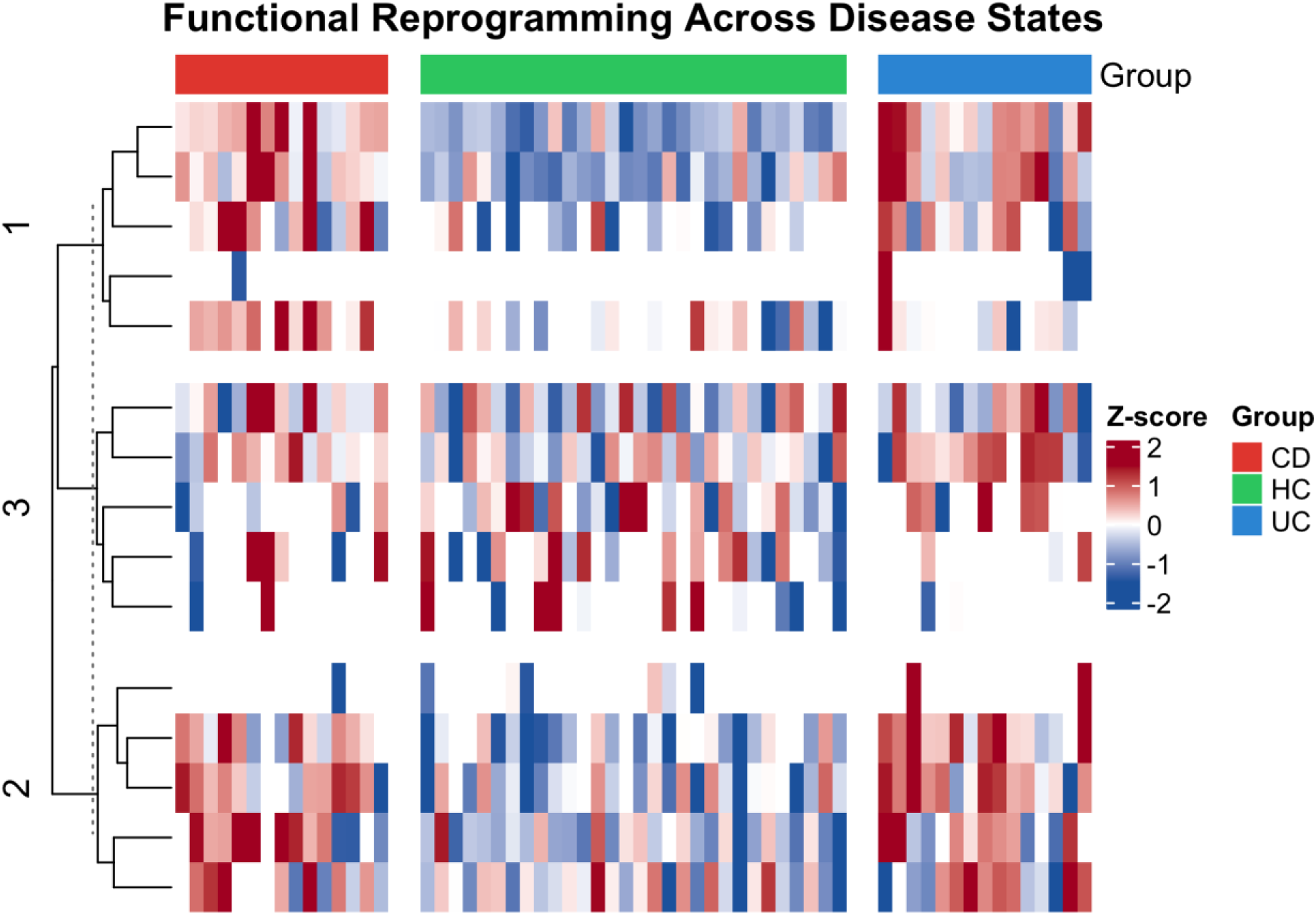
Heatmap of functional pathway abundance across HC, CD, and UC samples. Rows represent pathways and columns represent individual samples. Hierarchical clustering reveals partial grouping of samples; however, no clear separation between disease and healthy conditions is observed.

These results suggest that pathway-level variation alone does not provide strong discriminatory power for disease classification, consistent with previous reports that functional profiles remain relatively stable despite disease-associated dysbiosis.

These results demonstrate that pathway abundance patterns alone provide limited discriminatory power, reinforcing the need to examine higher-order structural organization within the microbiome. This lack of clear separation suggests that disease-associated alterations are not primarily encoded at the level of pathway abundance, but rather emerge from changes in interaction structure and network organization.

### 4.3 GLOBAL FUNCTIONAL ORGANIZATION (BETA DIVERSITY)

To assess global differences in microbiome functional organization across conditions, we performed beta diversity analysis based on pathway abundance profiles. This approach evaluates overall similarity and dissimilarity between samples, providing a system-level view of functional variation.

Beta diversity analysis revealed limited global separation between healthy controls and disease groups, and these differences were not statistically significant (PERMANOVA, p > 0.05). Although minor shifts in sample distribution were observed, CD, UC, and HC samples largely overlapped in ordination space, indicating that overall functional composition remains broadly similar across conditions.

The absence of strong clustering suggests that disease-associated changes are not primarily driven by large-scale differences in pathway presence or abundance. Instead, these findings point toward more subtle alterations that are not captured by global compositional metrics.

**Figure 3.**
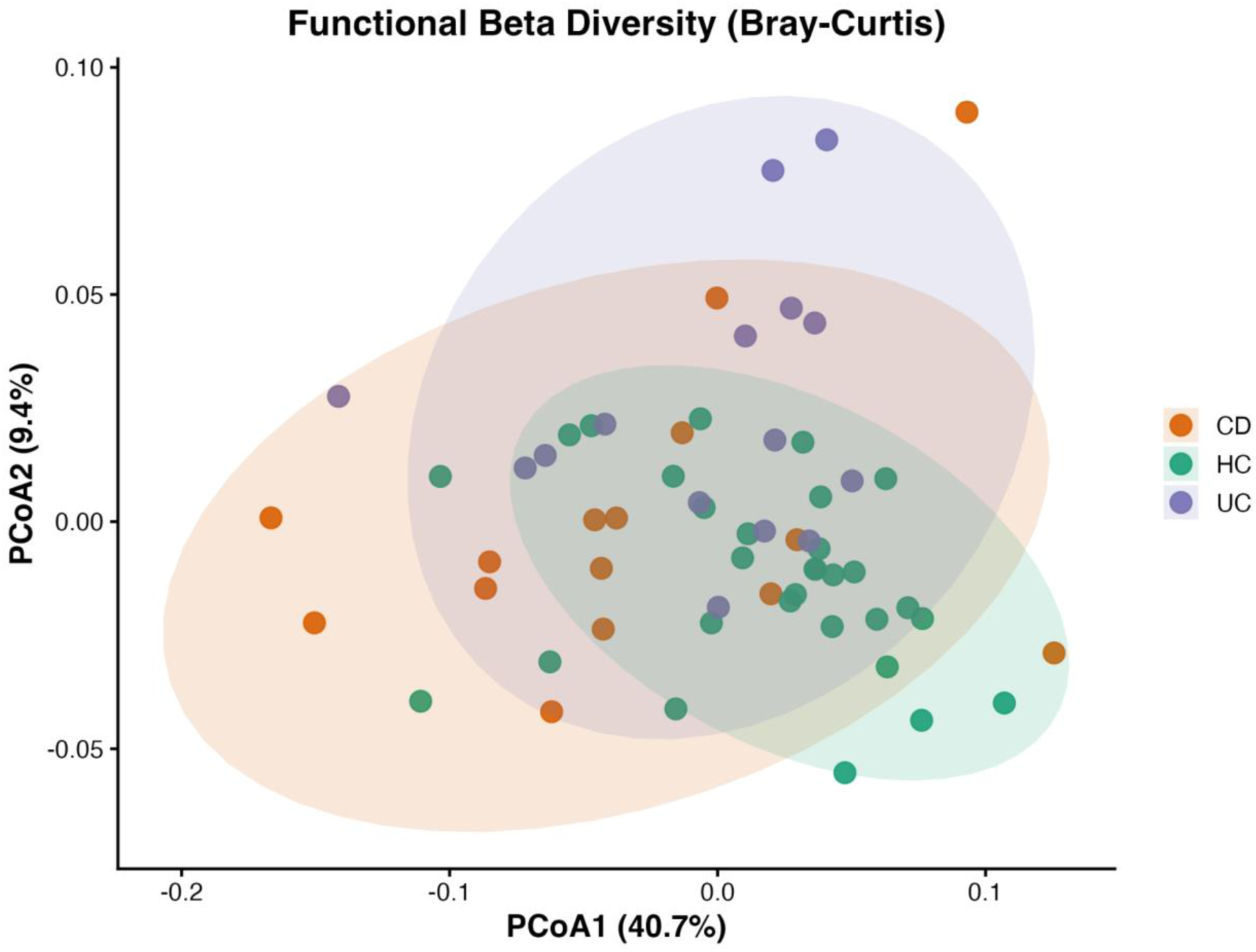
Beta diversity analysis of functional pathway profiles across HC, CD, and UC samples. Ordination plots show substantial overlap between groups, indicating limited global separation based on pathway abundance.

These observations reinforce that global compositional similarity masks deeper system-level differences, indicating that microbiome dysfunction is driven by alterations in interaction architecture rather than overall pathway presence.

### 4.4 FUNCTIONAL NETWORK REORGANIZATION

To investigate whether disease-associated changes are reflected at the level of functional interactions, we constructed correlation-based networks of microbial pathways for each group. This approach captures higher-order relationships between pathways, enabling analysis of system-level organization beyond individual pathway abundance.

Network analysis revealed substantial differences in functional organization between healthy and disease states. Healthy control networks exhibited higher connectivity and more distributed topology, characterized by increased network density and inter-module connectivity.

In contrast, disease networks showed reduced global connectivity and increased modular fragmentation, reflected by lower network density and higher modularity (Figure 4). CD networks displayed localized clustering, while UC networks exhibited stronger fragmentation and reduced inter-module interactions.

**Figure 4.**
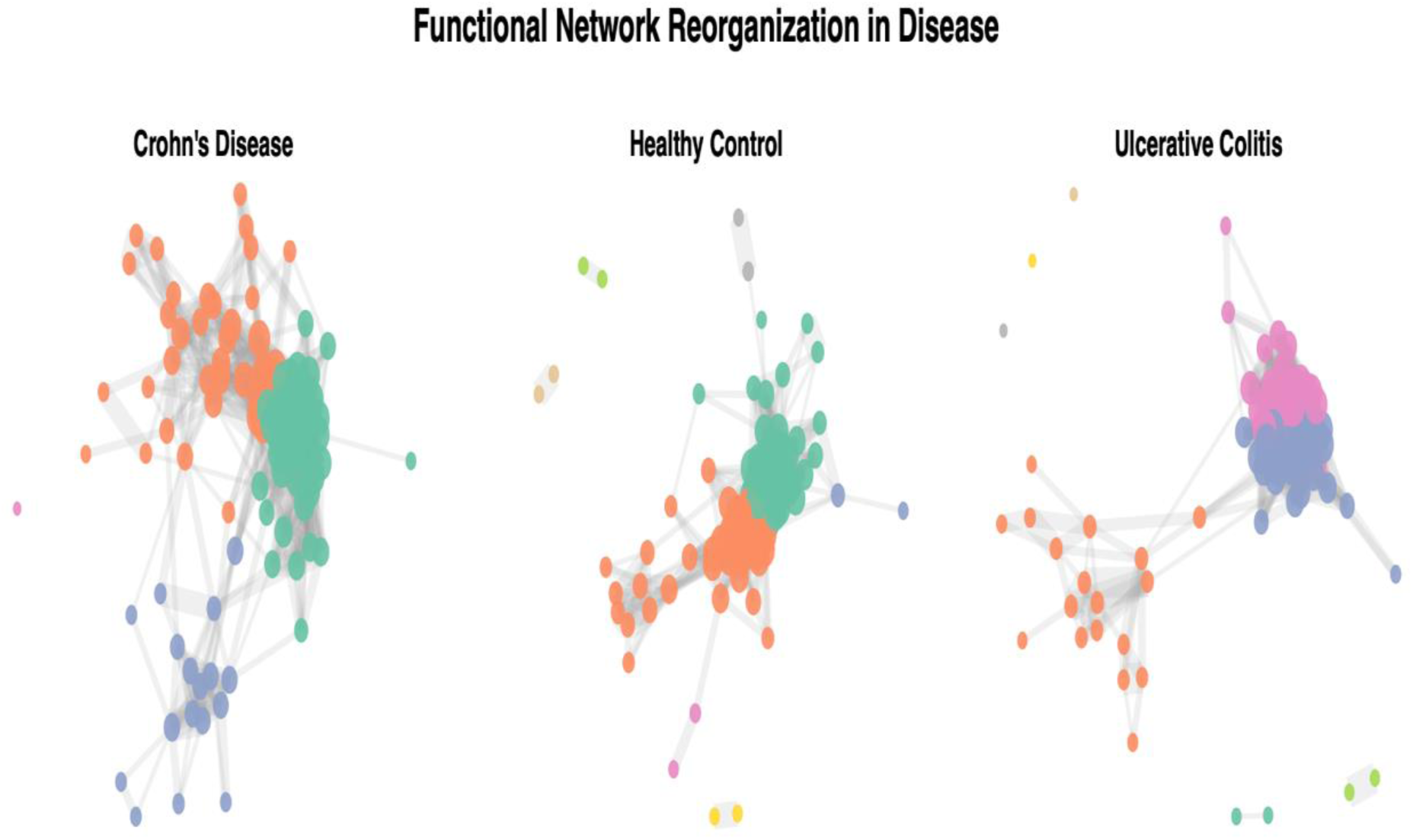
Functional network organization across HC, CD, and UC microbiomes. Nodes represent functional pathways and edges represent significant associations. Healthy networks exhibit distributed connectivity, whereas disease networks show increased clustering and fragmentation, indicating structural reorganization.

These findings indicate a transition from integrated to compartmentalized network architecture in disease conditions. Notably, these structural alterations were not captured by abundance-based analyses, highlighting the importance of interaction-level investigation.

These findings demonstrate that microbiome dysfunction in IBD is characterized by large-scale reorganization of functional interactions rather than changes in pathway abundance alone. This structural reorganization reflects a shift from globally integrated networks to compartmentalized systems, reducing coordination between pathways and potentially limiting adaptive responses under stress conditions.

### 4.5 KEYSTONE PATHWAY ARCHITECTURE

To identify key functional pathways driving microbiome network organization, differential betweenness centrality analysis was performed across disease comparisons (CD vs HC, UC vs HC, and CD vs UC).

Differential centrality analysis revealed substantial reorganization of keystone pathways across disease states. In both CD and UC, pathways associated with amino acid metabolism, nucleotide biosynthesis, and energy metabolism exhibited increased centrality, indicating their elevated role in maintaining network connectivity. In contrast, pathways related to carbohydrate metabolism and degradation processes showed reduced centrality, suggesting loss of distributed functional redundancy.

Notably, CD vs HC comparisons demonstrated stronger shifts in centrality compared to UC, indicating a more pronounced network restructuring in Crohn’s disease. The CD vs UC comparison further highlighted distinct pathway-level differences, supporting disease-specific functional architectures.

**Figure 5.**
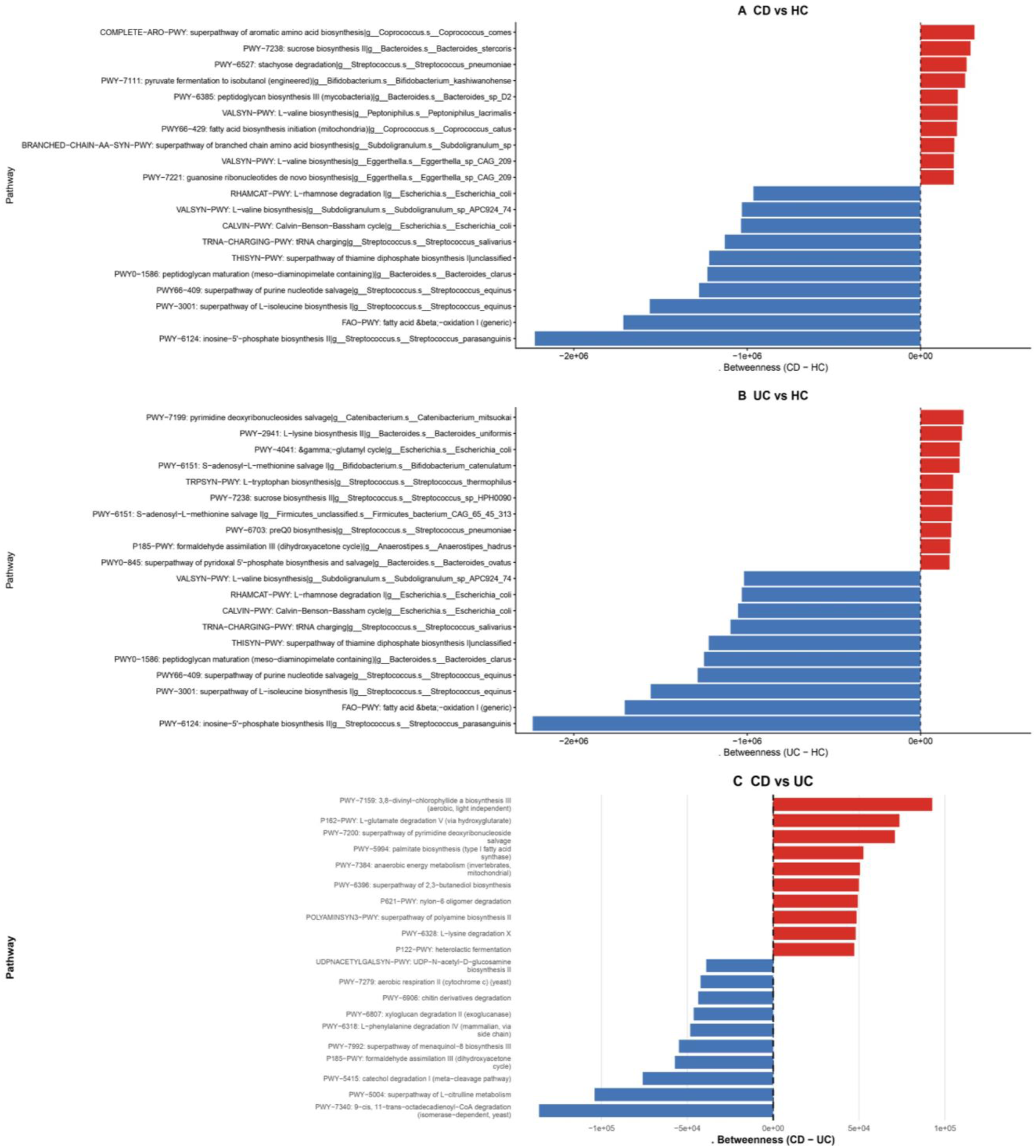
Differential keystone pathway analysis across disease conditions. (A) CD vs HC, (B) UC vs HC, and (C) CD vs UC comparisons showing pathways with increased (red) and decreased (blue) betweenness centrality. Positive values indicate pathways gaining centrality, while negative values indicate loss of centrality.

These findings indicate that disease progression is associated with a shift from distributed metabolic functionality toward centralized keystone pathways, reflecting increased network fragility and dependency on specific functions. This shift toward centralized keystone pathways indicates a transition from distributed functional control to bottleneck-driven regulation, where a limited subset of pathways disproportionately governs network stability, thereby increasing system fragility in disease states.

### 4.6 NETWORK ROBUSTNESS AND STABILITY

To evaluate the resilience of microbial functional networks, we assessed network robustness under simulated perturbations. Network robustness reflects the ability of a system to maintain connectivity and functional integrity when key components are disrupted.

Robustness analysis revealed clear differences in stability between healthy and disease networks. Under simulated node removal, HC networks maintained higher levels of connectivity, indicating strong resilience to perturbation.

In contrast, both CD and UC networks showed rapid loss of connectivity following node removal, demonstrating increased fragility. Targeted removal of highly connected nodes further amplified this effect, particularly in disease networks, where network integrity declined sharply.

Quantitative assessment using area under the curve (AUC) demonstrated reduced robustness in disease networks compared to HC, and this difference was statistically significant (p < 0.05, permutation test). These findings indicate that disease-associated network reorganization results in structurally weaker and less resilient systems.

**Figure 6.**
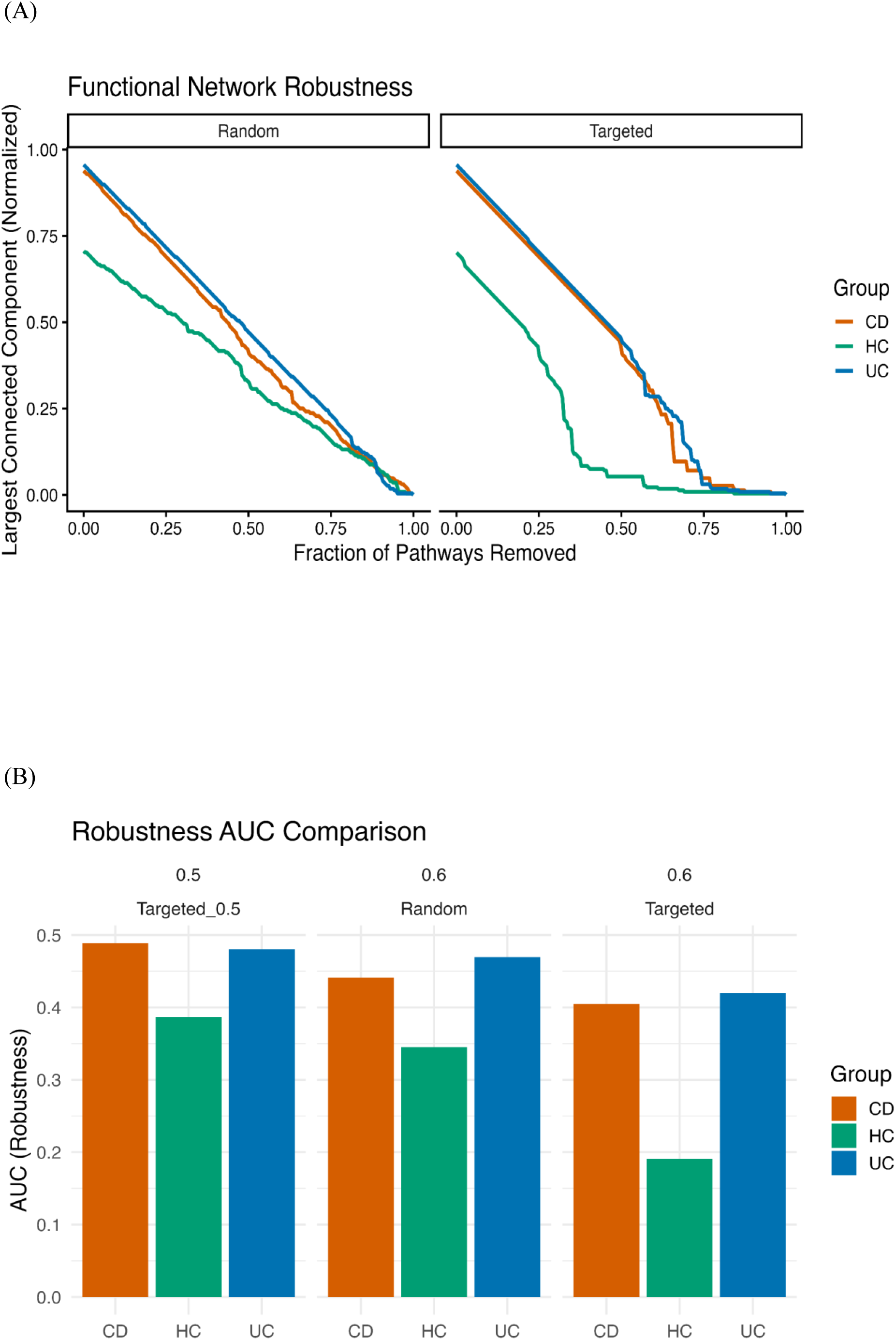
Functional network robustness analysis. (A) Robustness curves showing the decline in the largest connected component under random and targeted removal of pathways. Disease networks exhibit faster fragmentation, indicating reduced robustness compared to healthy controls. (B) AUC comparison: summarizing robustness across groups, confirming decreased stability in disease conditions.

These findings indicate that disease-associated microbiome networks are structurally fragile and operate under reduced tolerance to perturbation. The rapid loss of connectivity observed in disease conditions reflects a transition toward unstable system configurations, where network integrity depends on a limited set of highly connected pathways. This fragility is consistent with the observed centralization of network architecture, suggesting that disease-associated reorganization leads to increased vulnerability and reduced resilience at the systems level.

### 4.7 INTEGRATIVE NETWORK - REDUNDANCY ANALYSIS

To investigate the relationship between functional redundancy and network structure, we performed integrative analysis by correlating redundancy scores with network centrality metrics. This approach enables assessment of whether highly redundant pathways also occupy structurally important positions within the network.

Correlation analysis revealed weak and non-significant associations between redundancy and centrality metrics (Spearman ρ ≈ 0, p > 0.05), indicating decoupling between functional redundancy and structural importance. In HC, redundancy showed limited association with centrality, suggesting that highly connected pathways are not necessarily the most redundant. In disease states, this relationship became further decoupled. Both CD and UC networks exhibited increased variability, with no consistent alignment between redundancy and structural importance. In several cases, pathways with low redundancy displayed high centrality, indicating disproportionate influence despite limited functional backup.

These findings demonstrate a clear disconnect between redundancy and network structure, suggesting that functional importance within microbial systems is governed by interaction architecture rather than redundancy alone.

**Figure 7.**
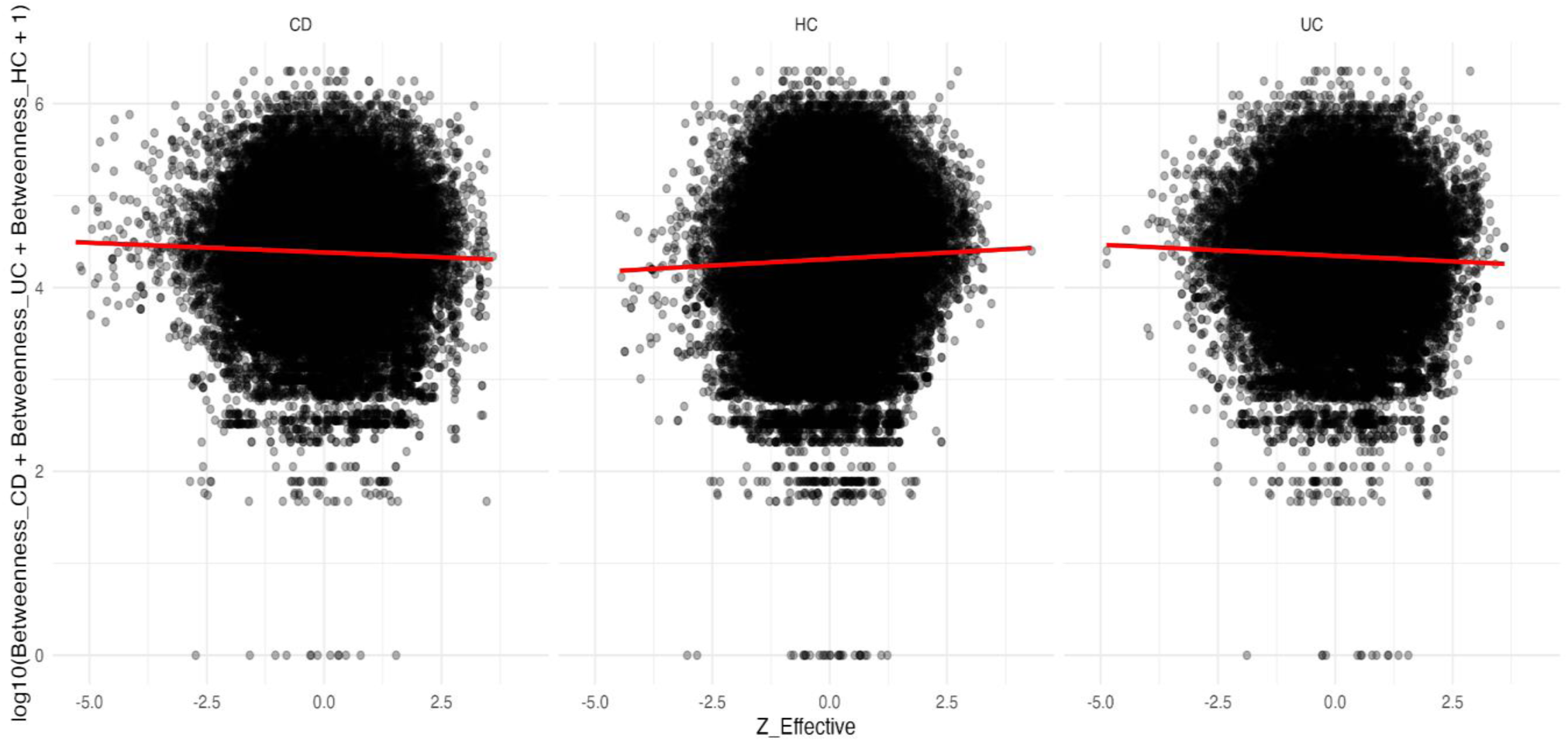
Relationship between functional redundancy and network centrality across HC, CD, and UC groups. Scatter plots show weak and inconsistent correlations between redundancy scores and betweenness centrality, indicating decoupling of redundancy and structural importance.

These results indicate that redundancy does not reliably predict structural importance, reinforcing the concept that microbiome function is governed by network organization rather than pathway redundancy alone.

### 4.8 MACHINE LEARNING VALIDATION OF FUNCTIONAL ARCHITECTURE

To evaluate whether structural features of microbiome functional organization can discriminate between disease states, we applied multiple machine learning models, including logistic regression, random forest, and SVM. This analysis provides predictive validation of whether network-derived features capture biologically meaningful disease signals. All models were evaluated using 5-fold cross-validation, and performance metrics were averaged across folds to ensure robustness and minimize overfitting.

#### 4.8.1 Model Performance Comparison

All models demonstrated consistent classification performance across disease groups, with similar trends observed across cross-validation folds. Logistic regression achieved the highest accuracy (0.824), followed by random forest (0.765) and SVM (0.706), indicating that disease-associated patterns are detectable across both linear and nonlinear frameworks.

**Figure 8.**
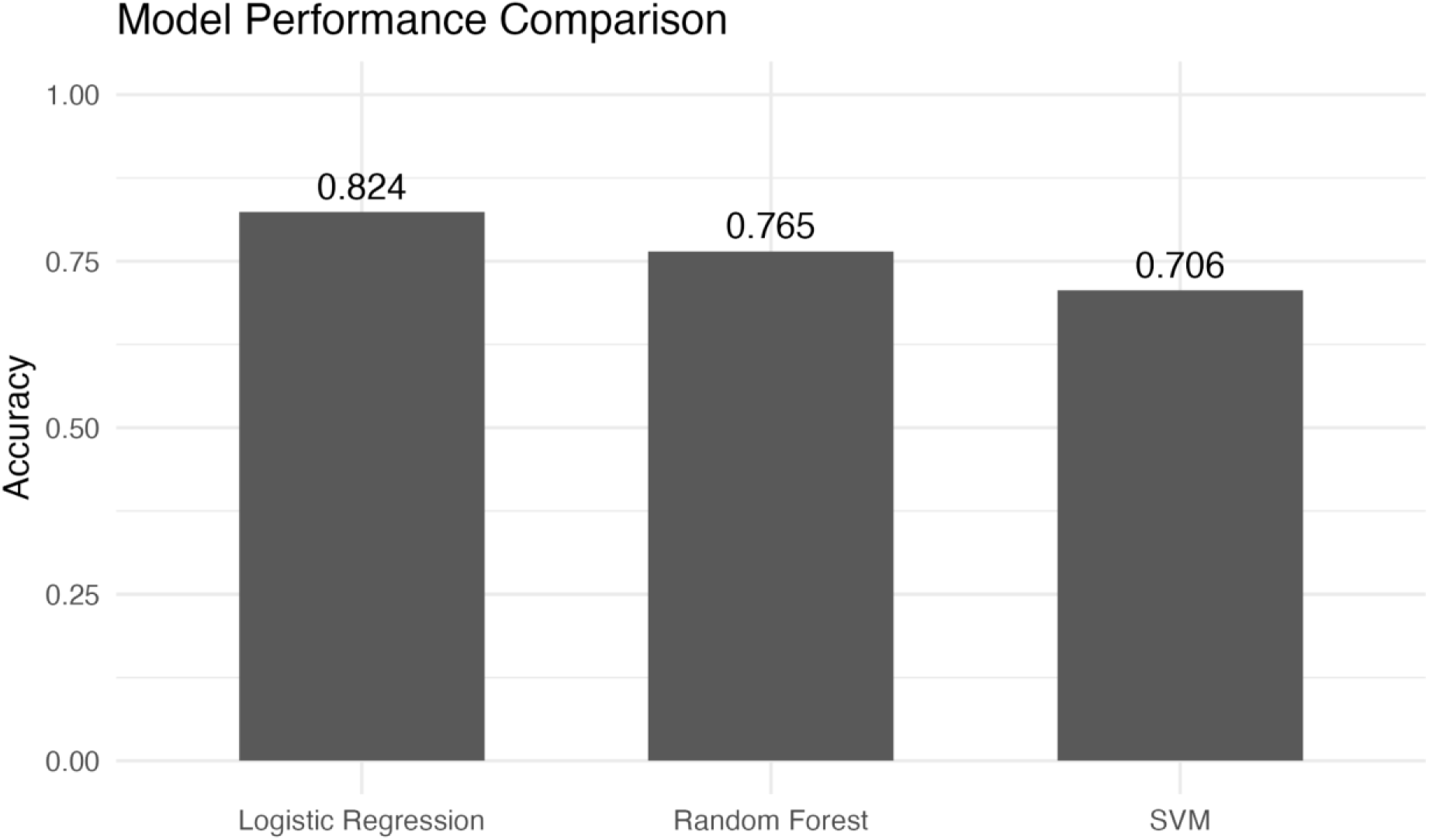
Comparison of classification accuracy across machine learning models. Logistic regression achieved the highest accuracy, followed by random forest and SVM.

These results suggest that disease-associated functional changes are structured and separable and do not require highly complex nonlinear decision boundaries for classification.

#### 4.8.2 Classification Behavior (Confusion Matrix)

To further assess classification performance, we examined the confusion matrix for the random forest model.

**Figure 9.**
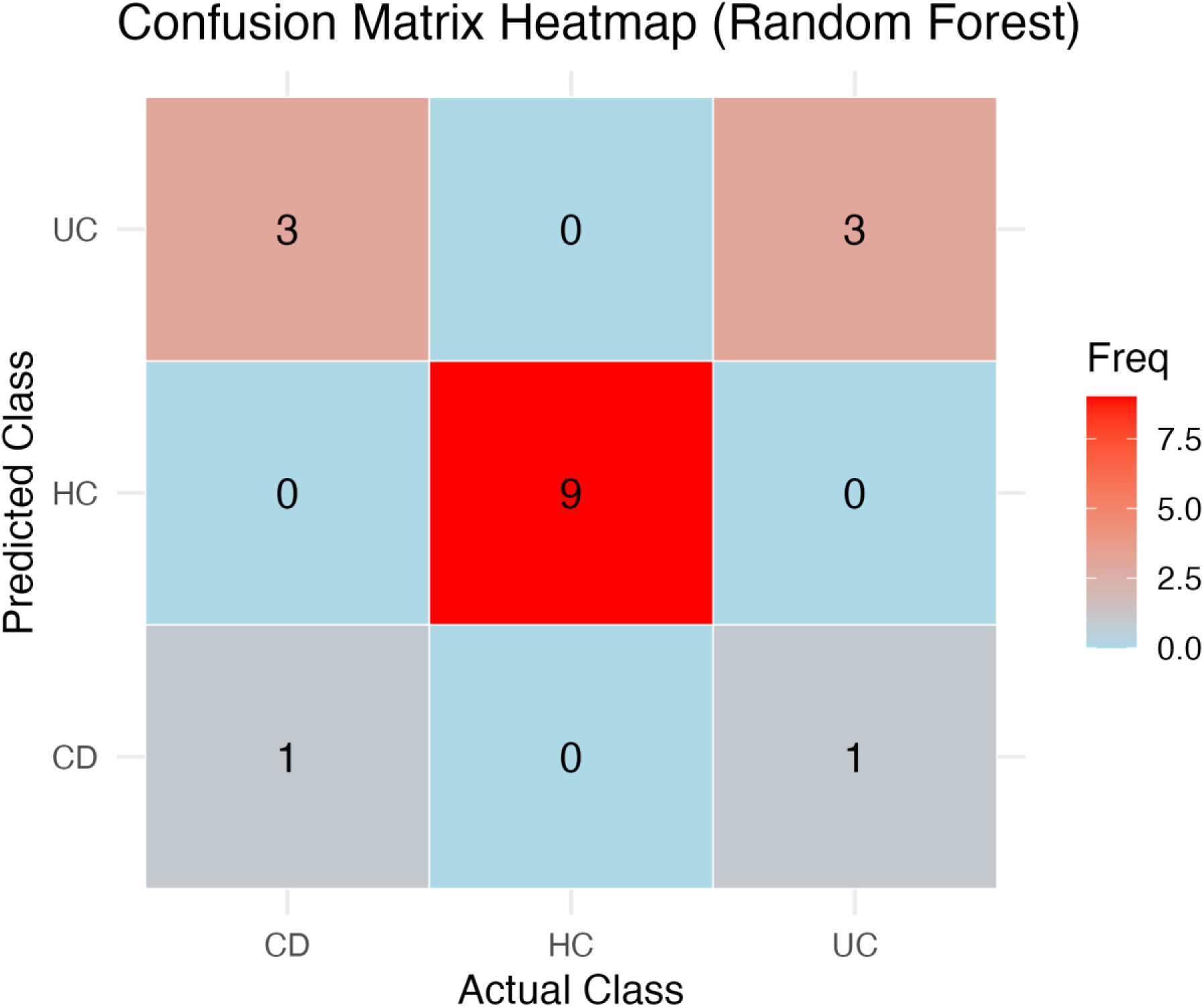
Confusion matrix for random forest classification across CD, UC, and HC groups.

The confusion matrix revealed that HC samples were classified with high accuracy, whereas misclassification occurred primarily between CD and UC groups. This indicates that disease states share partially overlapping functional characteristics, while remaining clearly distinguishable from healthy microbiome profiles.

#### 4.8.3 Model Discrimination (ROC Analysis)

ROC curves were used to evaluate model discrimination across classes.

**Figure 10.**
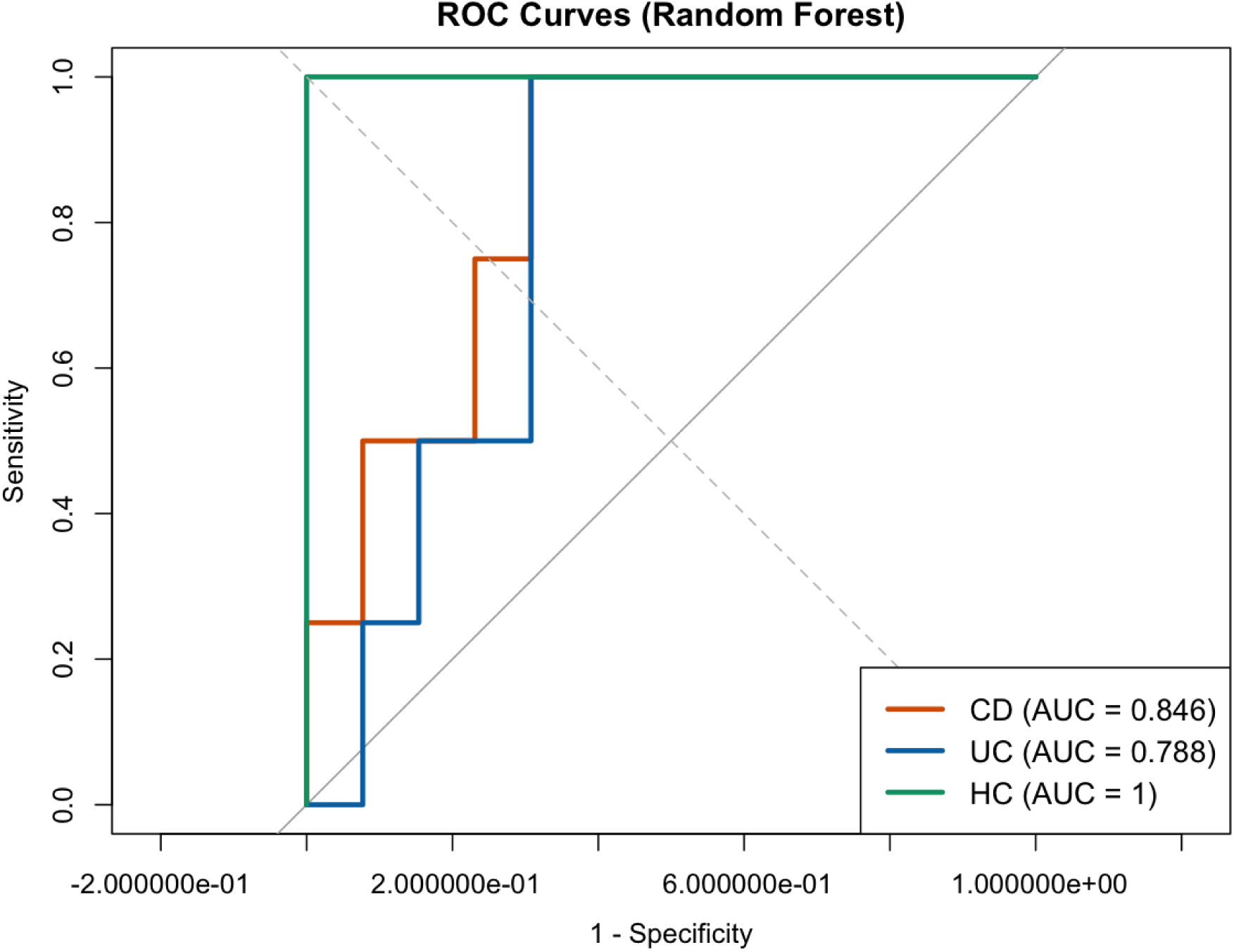
ROC curves for random forest classification across CD, UC, and HC groups. AUC values indicate classification performance.

ROC analysis confirmed strong model performance, with high AUC values observed across all classes. Notably, healthy controls exhibited near-perfect classification, while CD and UC showed moderate separation. These findings reinforce that disease conditions retain partial functional similarity yet remain distinguishable at the systems level.

#### 4.8.4 Feature Importance and Predictive Drivers

To identify key features driving classification performance, feature importance was assessed using the random forest model.

**Figure 11.**
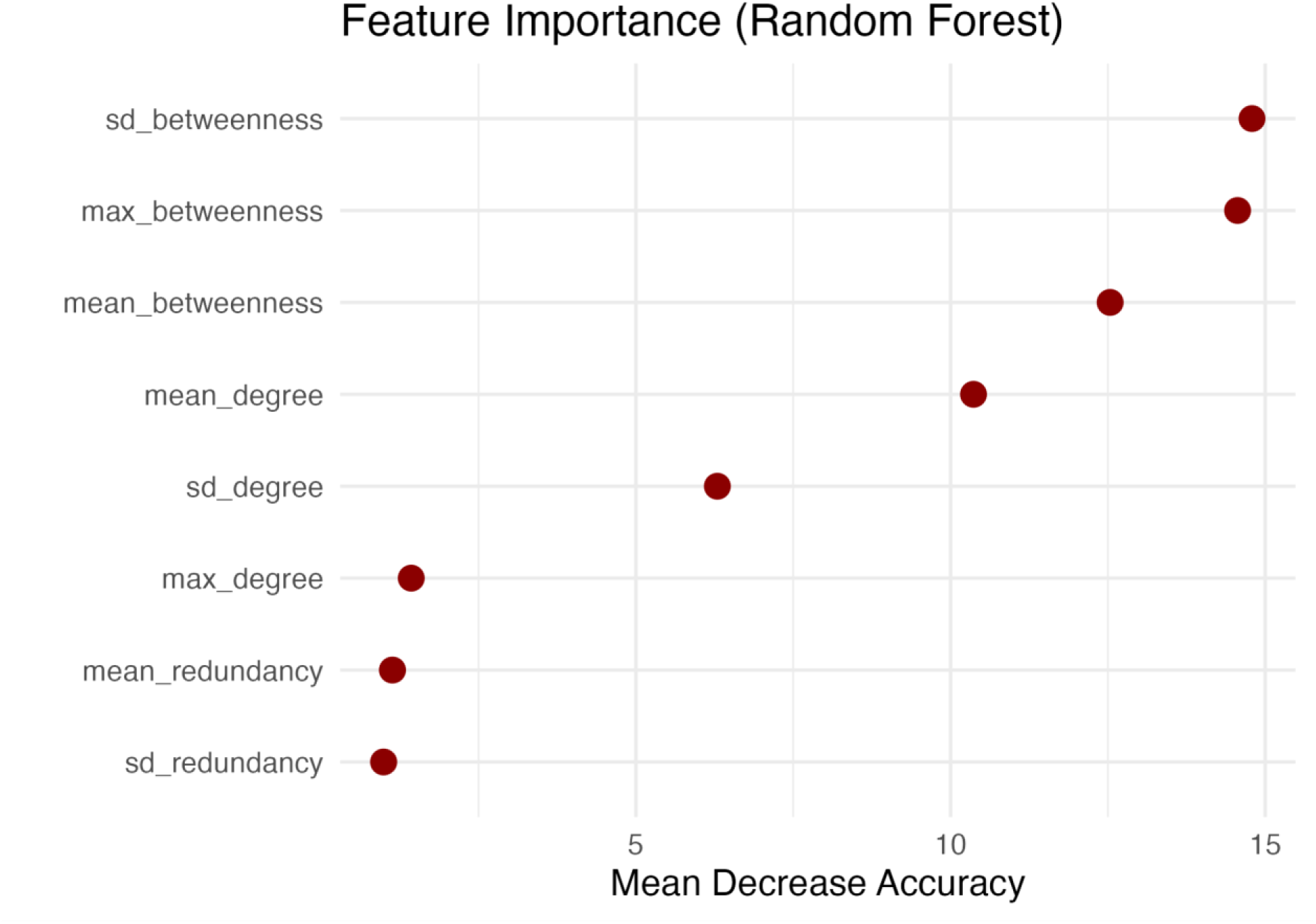
Feature importance derived from Random Forest classification. Features are ranked based on mean decrease in accuracy.

Feature importance analysis revealed that network-derived features, particularly betweenness centrality (mean, maximum, and variability), were the strongest predictors of disease states, whereas redundancy-based features contributed minimally. Collectively, these results provide strong predictive validation that microbiome dysfunction in disease is primarily driven by structural reorganization rather than loss of functional redundancy.

## DISCUSSION

The present study provides a systems-level characterization of microbiome functional organization in IBD, demonstrating that disease-associated alterations are primarily encoded in the architecture of functional interactions rather than in the loss of functional capacity. By integrating redundancy profiling, network topology, keystone pathway analysis, robustness assessment, and machine learning validation, we show that microbiome dysfunction in IBD reflects a coordinated reorganization of functional relationships, consistent with emerging systems-level perspectives of microbial ecosystems [8,18,21].

Consistent with initial compositional analyses, redundancy and pathway abundance exhibited limited discriminatory power between HC and disease states. Functional redundancy differences were modest, and both pathway composition and beta diversity analyses revealed substantial overlap across groups. These findings indicate that the overall functional repertoire of the microbiome remains broadly preserved in IBD, supporting previous observations of functional stability despite taxonomic shifts [6,8,29]. However, this apparent stability masks deeper structural alterations in how these functions are organized and interact.

In contrast, network-level analyses revealed pronounced differences in system organization. Healthy microbiomes exhibited a distributed and highly connected architecture, consistent with coordinated and resilient functional interactions. Disease networks, however, showed increased modular fragmentation and localized clustering, indicating a transition toward compartmentalized organization. This shift is consistent with prior studies demonstrating disruption of microbial interaction networks in disease states [18,19,21].

The reconfiguration of keystone pathway architecture further reinforces this transition. In healthy states, keystone pathways were distributed across the network, reflecting decentralized control and functional redundancy in regulatory influence. In disease conditions, keystone pathways became increasingly concentrated, indicating a shift toward centralized control. This redistribution aligns with ecological network theory, where centralization increases system vulnerability and dependency on key nodes [10,25].

These structural changes have direct consequences for network stability. Robustness analysis demonstrated that disease networks are significantly more fragile, exhibiting rapid loss of connectivity under perturbation. In contrast, healthy networks maintained structural integrity, reflecting inherent resilience. This observation is consistent with studies linking network fragmentation to reduced ecosystem stability and resilience [26].

Importantly, integrative analysis revealed a clear decoupling between functional redundancy and network centrality. Pathways with high structural importance were not necessarily those with high redundancy, indicating that redundancy does not reliably predict functional influence within the system. This finding challenges traditional assumptions in microbiome research and supports recent work suggesting that functional redundancy alone cannot explain ecosystem stability [22,23,29].

Machine learning analysis provided independent and predictive validation of these observations. Models trained on network-derived features consistently outperformed those based on redundancy metrics, with centrality measures, particularly betweenness-related features, emerging as the strongest predictors of disease state. This aligns with recent studies highlighting the predictive power of network-based features in microbiome classification tasks [14,28].

Notably, while previous studies have independently examined functional redundancy or microbial network structure [1,8,21,22,23], this study uniquely integrates redundancy, network topology, robustness, and machine learning within a single analytical framework. This integrative approach enables direct comparison between compositional and structural determinants of microbiome dysfunction, providing evidence that interaction architecture, rather than functional presence alone, is the dominant driver of disease-associated changes.

Collectively, these findings support a revised conceptual framework for microbiome dysfunction in IBD. Rather than being driven by loss of functional components, disease is characterized by a transition from distributed, redundant, and resilient networks to centralized, fragmented, and fragile architectures. This systems-level shift is consistent with principles of complex systems theory, where loss of network integration precedes functional collapse [21,26].

Despite these advances, several limitations should be acknowledged. The networks analyzed in this study are correlation-based and may not fully capture causal or directional relationships between pathways. Additionally, correlation-based network inference may introduce spurious associations, a known limitation in microbiome network analysis [19,20]. Future work integrating longitudinal sampling, causal inference methods, and experimental validation will be essential to further resolve the mechanistic basis of microbiome network reorganization. Furthermore, the cross-sectional design of this study limits the ability to capture temporal dynamics of microbiome reorganization during disease progression.

In conclusion, this study demonstrates that microbiome dysfunction in IBD is fundamentally a problem of organization rather than composition. By demonstrating that disease-associated signals are embedded in network structure rather than functional abundance, this study establishes a systems-level framework for understanding microbiome-driven disease. These findings suggest that future therapeutic strategies should focus not only on restoring microbial composition but also on re-establishing functional interaction networks to enhance system stability and resilience.

## CONCLUSION

In this study, we present a comprehensive systems-level analysis of microbiome functional organization in IBD [8,21,22], demonstrating that disease-associated alterations are primarily encoded in the structure of functional interactions rather than in the loss of functional capacity. By integrating pathway composition, network topology, keystone pathway analysis, robustness assessment, and machine learning validation, we show that microbiome dysfunction reflects a coordinated reorganization of interaction architecture.

Despite substantial overlap in functional composition and redundancy across conditions, network-based analyses revealed pronounced structural differences. Disease states were characterized by increased modular fragmentation, centralized control of keystone pathways, and reduced network robustness, indicating a transition from distributed and resilient systems to fragile, bottlenecked architectures. These findings suggest that functional stability at the compositional level masks deeper organizational instability at the systems level.

Importantly, machine learning analysis provided independent validation of these observations, demonstrating that network-derived features particularly centrality metrics consistently outperform redundancy-based measures in predicting disease states. This confirms that structural properties of the microbiome encode more biologically relevant and diagnostically informative signals than traditional abundance- or redundancy-based approaches [14,28].

Collectively, our results support a revised framework in which microbiome dysfunction in IBD arises from reorganization of functional networks rather than depletion of functions. This shift in perspective emphasizes that disease is driven by changes in interaction architecture, leading to increased dependency on key pathways and reduced system resilience.

Overall, this study establishes that microbiome dysfunction in IBD is fundamentally a problem of network organization rather than functional loss. By identifying structural reorganization as the primary driver of disease-associated changes, this work provides a foundation for future therapeutic strategies aimed at restoring network stability and functional resilience.

## Supporting information

Supplementary Materials for IBD Functional Microbiome Analysis (Datasets, Figures, and Scripts)

## DATA AVAILABILITY

All data used in this study are publicly available from the NCBI Sequence Read Archive (SRA) under BioProject accession PRJNA945504. Processed datasets generated and analyzed during this study are provided as supplementary materials.

## CODE AVAILABILITY

Custom R scripts used for data preprocessing, network construction, statistical analysis, and machine learning are available from the corresponding author upon reasonable request.

## ETHICS STATEMENT

This study utilized publicly available, de-identified datasets from the NCBI Sequence Read Archive (SRA). No human participants or animals were directly involved; therefore, ethical approval and informed consent were not required.

## FUNDING

This research received no external funding.

## CONFLICT OF INTEREST

The authors declare no competing interests.

## AUTHOR CONTRIBUTIONS

Mihika Kenavdekar conceived and conducted the study, performed the analyses, interpreted the results, and drafted the manuscript. Dr. Elamathi Natarajan supervised the research, provided methodological guidance, and critically reviewed the manuscript. All authors approved the final version of the manuscript.

## ACKNOWLEDGMENTS

The authors acknowledge the NCBI Sequence Read Archive for providing open-access datasets used in this study. The authors also thank Biotecnika Info Labs Pvt. Ltd., Bengaluru, for providing computational support and research guidance.

## Notes

### Competing Interest Statement

The authors have declared no competing interest.

