## Supplementary Materials for IBD Functional Microbiome Analysis (Datasets, Figures, and Scripts) for "NETWORK-BASED FUNCTIONAL FRAGILITY REVEALS SYSTEM-LEVEL REORGANIZATION OF THE GUT MICROBIOME IN INFLAMMATORY BOWEL DISEASE": IBD_Functional_Network_Reorganization_Supplementary.pdf

### Figure S1: Differential Keystone Pathways

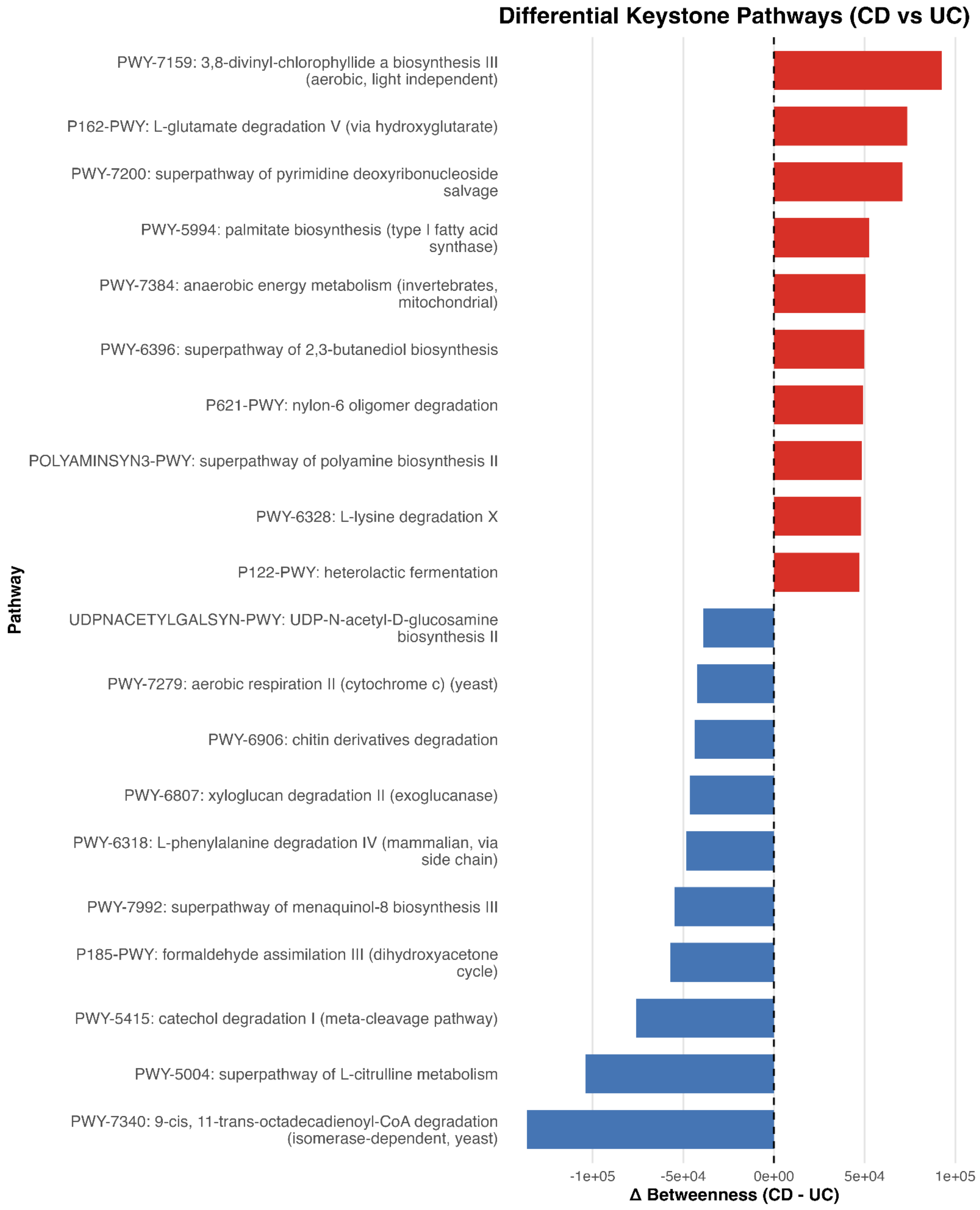

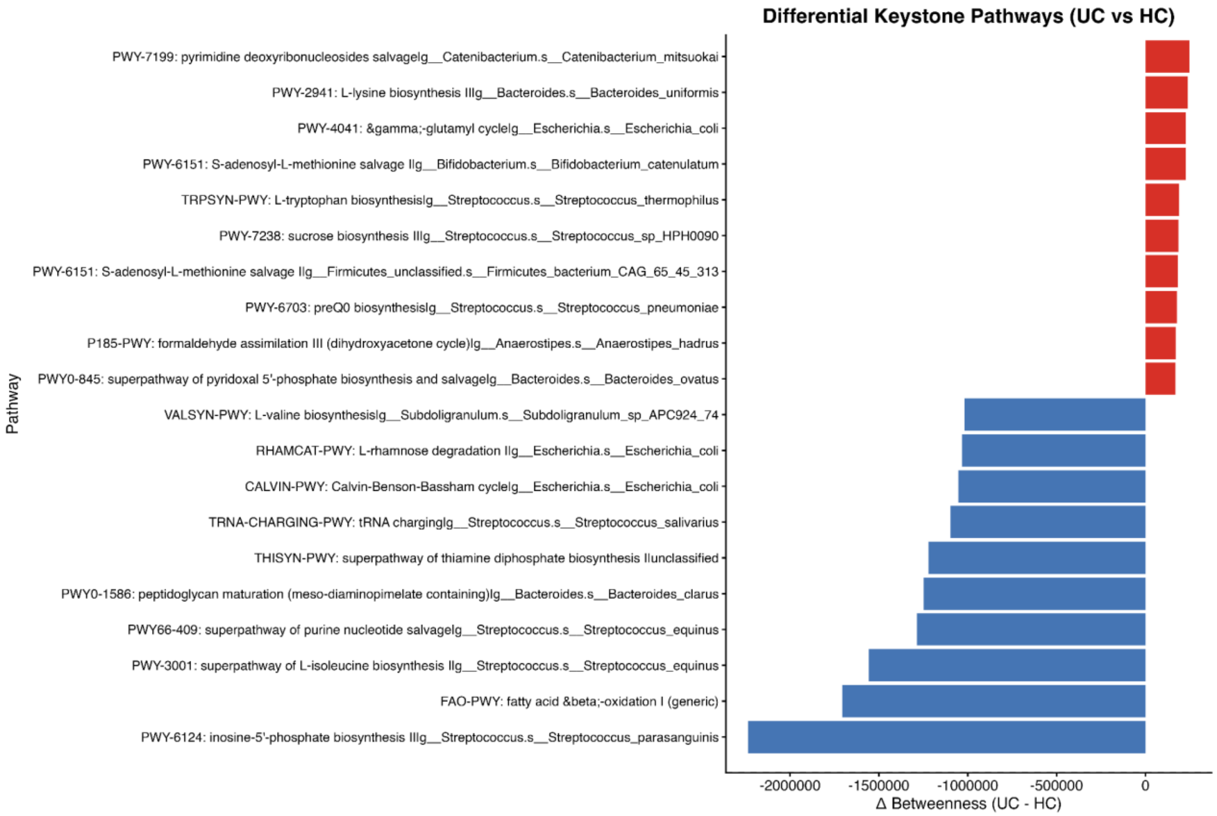
Differential Keystone Pathways (UC vs HC)
PWY-7199: pyrimidine deoxyribonucleosides salvagelg__Catenibacterium.s__Catenibacterium_mitsuokai
PWY-2941: L-lysine biosynthesis IIIg__Bacteroides.s__Bacteroides_uniformis
PWY-4041: γ-glutamyl cyclelg__Escherichia.s__Escherichia_coli
PWY-6151: S-adenosyl-L-methionine salvage IIg__Bifidobacterium.s__Bifidobacterium_catenulatum
TRPSYN-PWY: L-tryptophan biosynthesislg__Streptococcus.s__Streptococcus_thermophilus
PWY-7238: sucrose biosynthesis IIIg__Streptococcus.s__Streptococcus_sp_HPH0090
PWY-6151: S-adenosyl-L-methionine salvage IIg__Firmicutes_unclassified.s__Firmicutes_bacterium_CAG_65_45_313
PWY-6703: preQ0 biosynthesislg__Streptococcus.s__Streptococcus_pneumoniae
P185-PWY: formaldehyde assimilation III (dihydroxyacetone cycle)lg__Anaerostipes.s__Anaerostipes_hadrus
PWY0-845: superpathway of pyridoxal 5'-phosphate biosynthesis and salvagelg__Bacteroides.s__Bacteroides_ovatus
VALSYN-PWY: L-valine biosynthesislg__Subdoligranulum.s__Subdoligranulum_sp_APC924_74
RHAMCAT-PWY: L-rhamnose degradation IIg__Escherichia.s__Escherichia_coli
CALVIN-PWY: Calvin-Benson-Bassham cyclelg__Escherichia.s__Escherichia_coli
TRNA-CHARGING-PWY: tRNA charginglg__Streptococcus.s__Streptococcus_salivarius
THISYN-PWY: superpathway of thiamine diphosphate biosynthesis IIunclassified
PWY0-1586: peptidoglycan maturation (meso-diaminopimelate containing)lg__Bacteroides.s__Bacteroides_clarus
PWY66-409: superpathway of purine nucleotide salvagelg__Streptococcus.s__Streptococcus_equinus
PWY-3001: superpathway of L-isoleucine biosynthesis IIg__Streptococcus.s__Streptococcus_equinus
FAO-PWY: fatty acid β-oxidation I (generic)
PWY-6124: inosine-5'-phosphate biosynthesis IIIg__Streptococcus.s__Streptococcus_parasanguinis
Pathway
-2000000
-1500000
-1000000
-500000
0
Δ Betweenness (UC - HC)

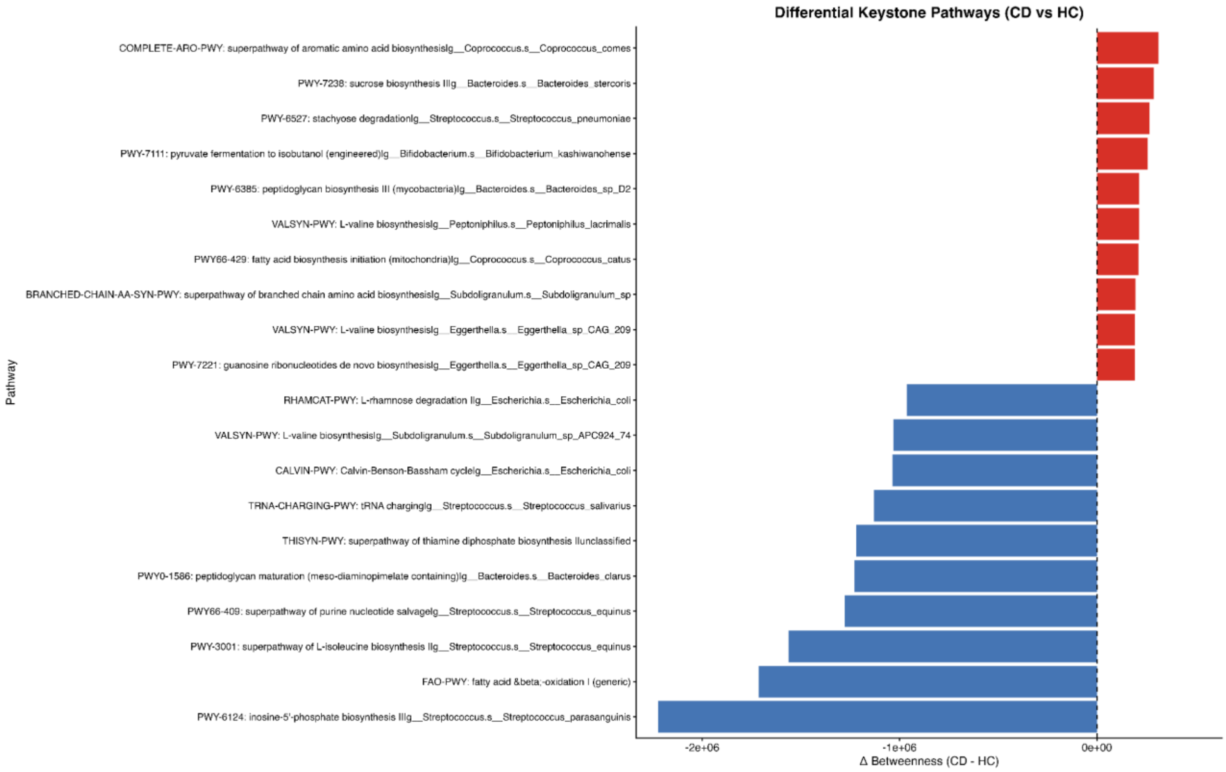
Differential Keystone Pathways (CD vs HC)
COMPLETE-ARO-PWY: superpathway of aromatic amino acid biosynthesislg__Coprococcus.s__Coprococcus_comes
PWY-7238: sucrose biosynthesis IIIg__Bacteroides.s__Bacteroides_stercoris
PWY-6527: stachyose degradationlg__Streptococcus.s__Streptococcus_pneumoniae
PWY-7111: pyruvate fermentation to isobutanol (engineered)lg__Bifidobacterium.s__Bifidobacterium_kashiwanohense
PWY-6385: peptidoglycan biosynthesis III (mycobacteria)lg__Bacteroides.s__Bacteroides_sp_D2
VALSYN-PWY: L-valine biosynthesislg__Peptoniphilus.s__Peptoniphilus_lacrimalis
PWY66-429: fatty acid biosynthesis initiation (mitochondria)lg__Coprococcus.s__Coprococcus_catus
BRANCHED-CHAIN-AA-SYN-PWY: superpathway of branched chain amino acid biosynthesislg__Subdoligranulum.s__Subdoligranulum_sp
VALSYN-PWY: L-valine biosynthesislg__Eggerthella.s__Eggerthella_sp_CAG_209
PWY-7221: guanosine ribonucleotides de novo biosynthesislg__Eggerthella.s__Eggerthella_sp_CAG_209
RHAMCAT-PWY: L-rhamnose degradation IIg__Escherichia.s__Escherichia_coli
VALSYN-PWY: L-valine biosynthesislg__Subdoligranulum.s__Subdoligranulum_sp_APC924_74
CALVIN-PWY: Calvin-Benson-Bassham cyclelg__Escherichia.s__Escherichia_coli
TRNA-CHARGING-PWY: tRNA charginglg__Streptococcus.s__Streptococcus_salivarius
THISYN-PWY: superpathway of thiamine diphosphate biosynthesis IIunclassified
PWY0-1586: peptidoglycan maturation (meso-diaminopimelate containing)lg__Bacteroides.s__Bacteroides_clarus
PWY66-409: superpathway of purine nucleotide salvagelg__Streptococcus.s__Streptococcus_equinus
PWY-3001: superpathway of L-isoleucine biosynthesis IIg__Streptococcus.s__Streptococcus_equinus
FAO-PWY: fatty acid β-oxidation I (generic)
PWY-6124: inosine-5'-phosphate biosynthesis IIIg__Streptococcus.s__Streptococcus_parasanguinis
Pathway
-2e+06
-1e+06
0e+00
Δ Betweenness (CD - HC)

Figure S1. Differential Keystone Pathways.
Bar plots illustrating changes in betweenness centrality (Δ Betweenness) across Crohn's Disease (CD), Ulcerative Colitis (UC), and Healthy Controls (HC). Positive values indicate pathways enriched in disease conditions, whereas negative values indicate pathways that are enriched in healthy controls.

**Figure S2: Redundancy vs Betweenness (All Groups)**

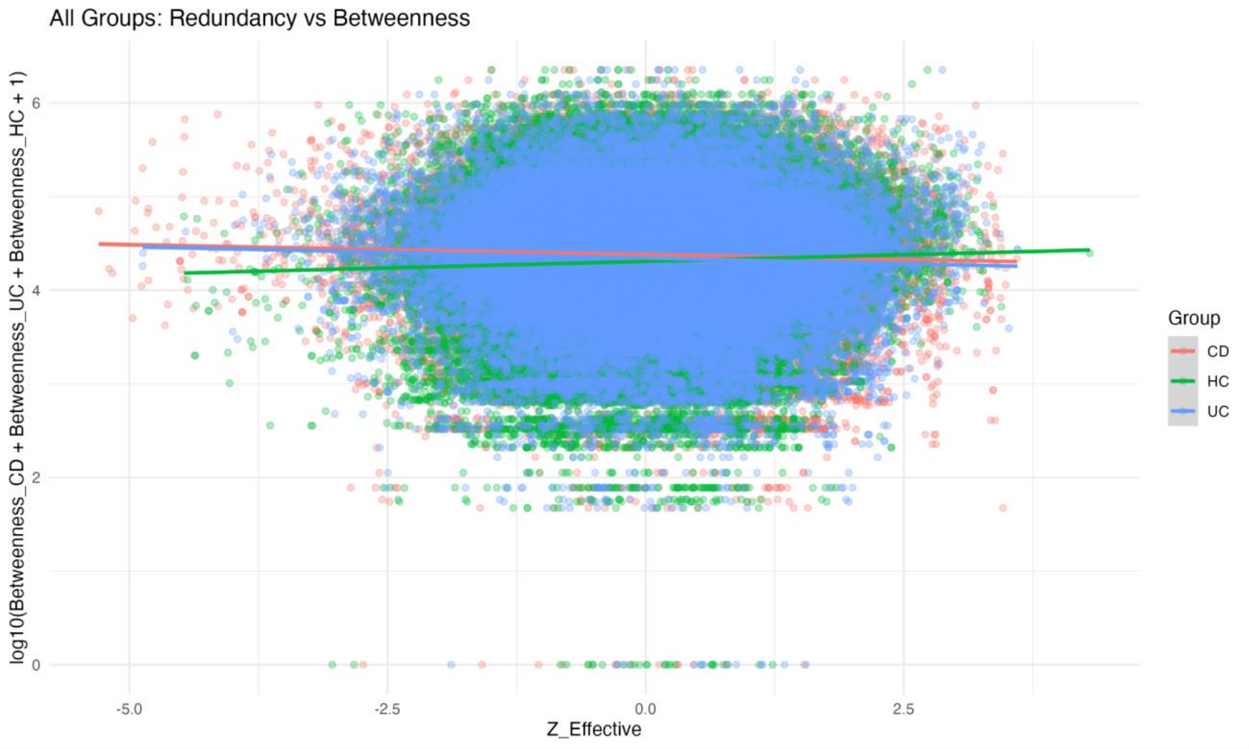


Figure S2. Functional Redundancy vs Betweenness Across All Groups.
Scatter plot showing the relationship between functional redundancy (Z-effective) and betweenness centrality across CD, UC, and HC samples, highlighting global network organization.

**Figure S3: Crohn's Disease Network Analysis**

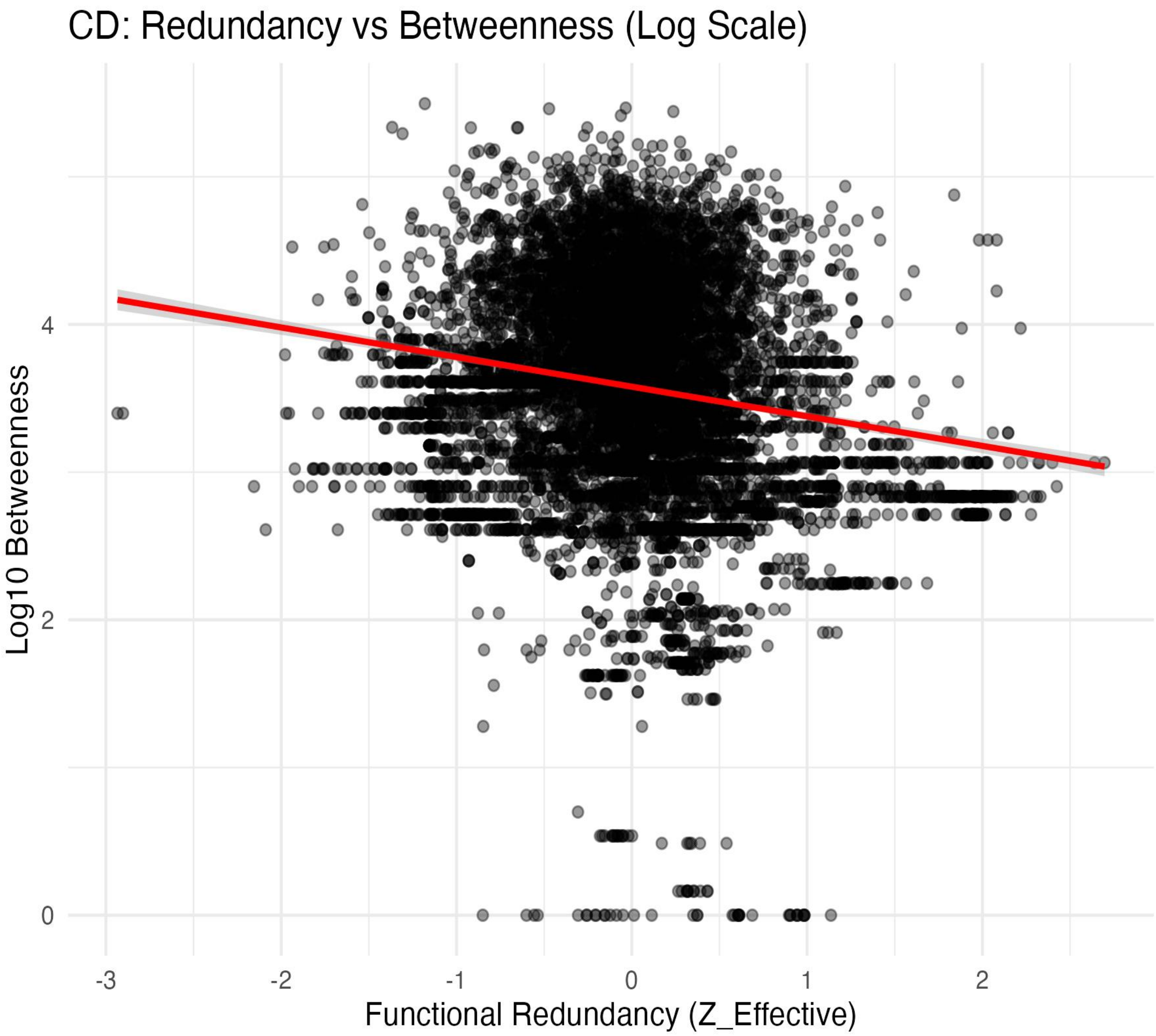

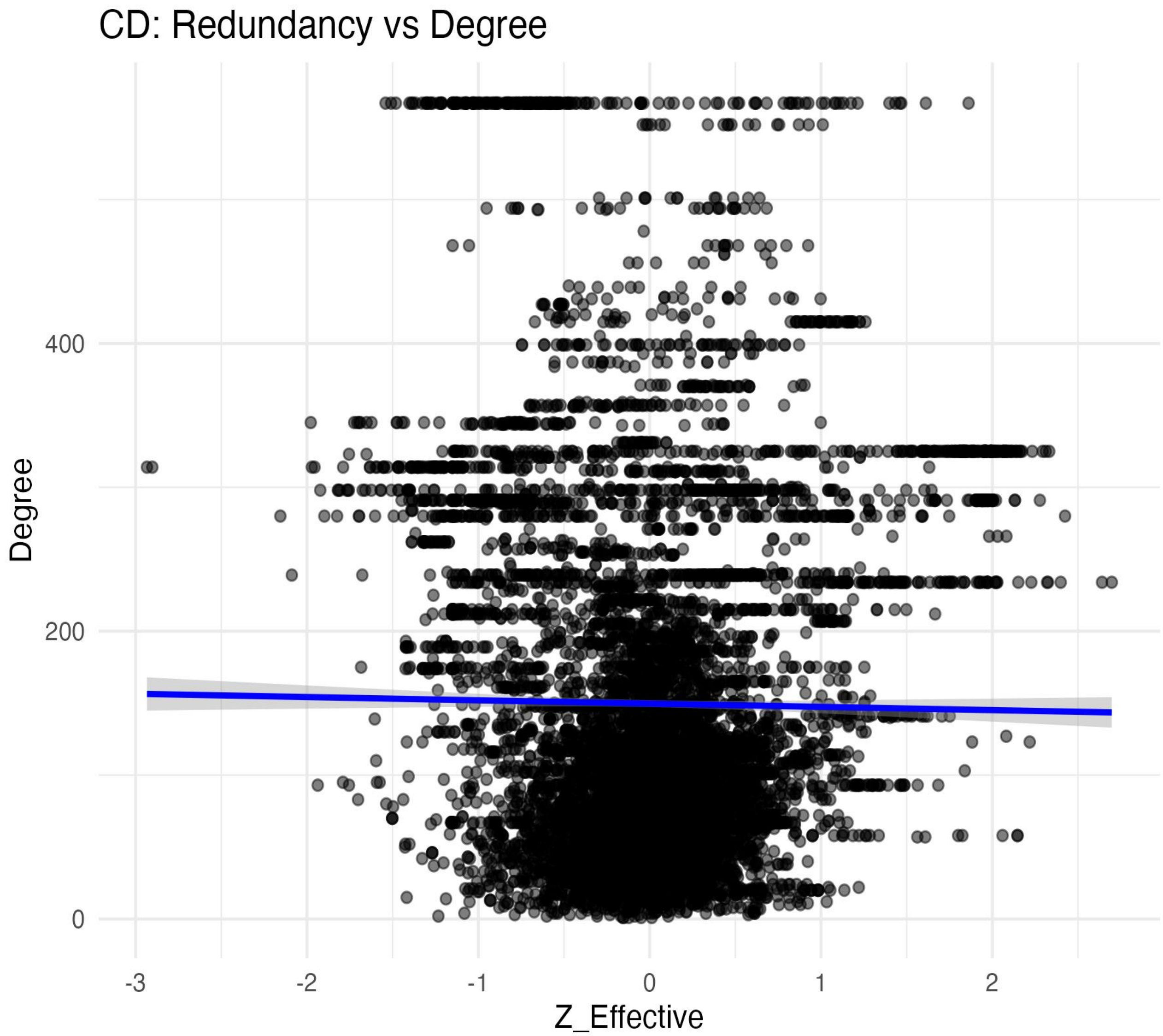


Figure S3. Network Topology in Crohn’s Disease.
Relationships between functional redundancy and network centrality metrics, illustrating structural alterations in CD microbiomes.

**Figure S4: Healthy Control Network Analysis**

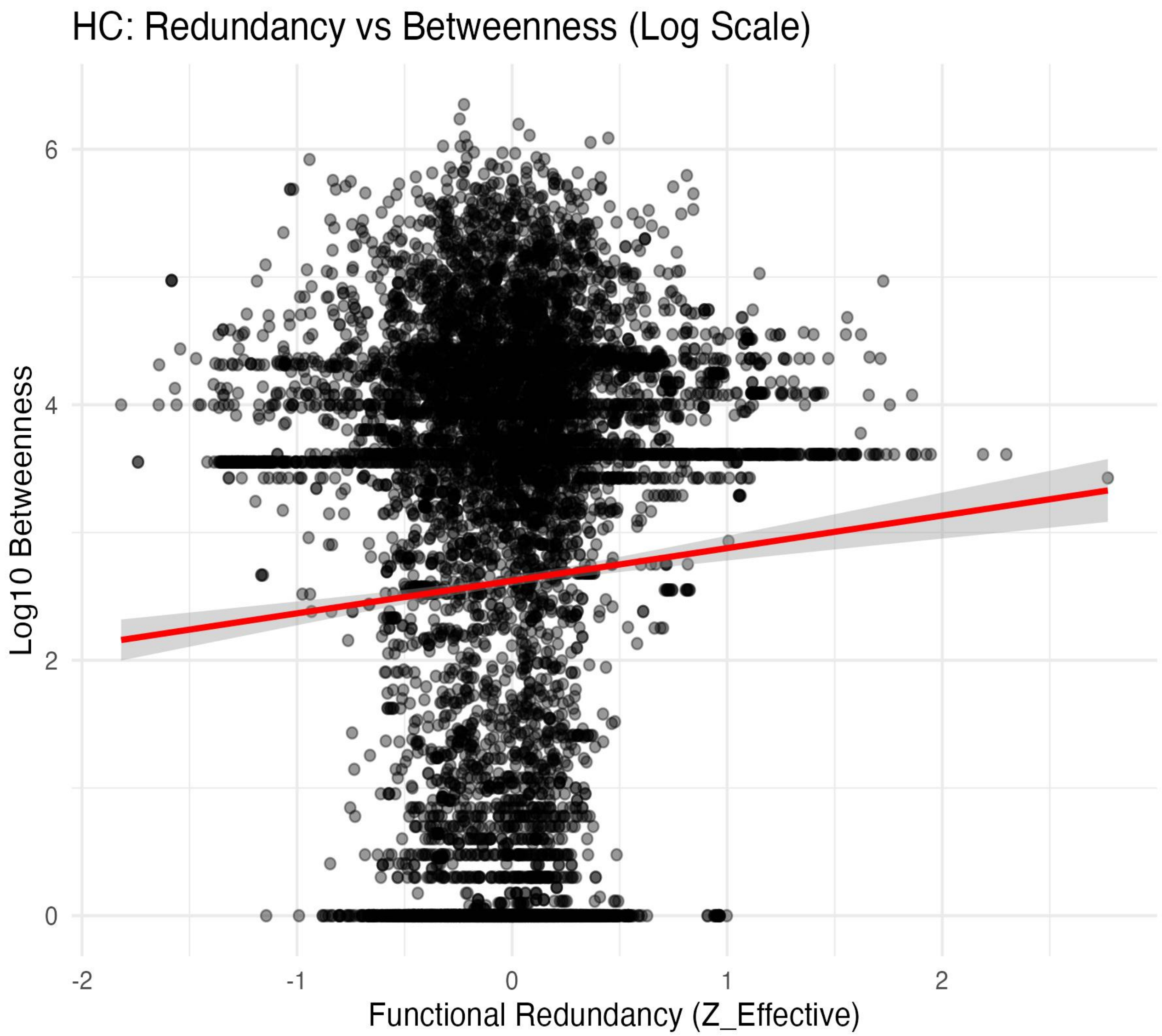

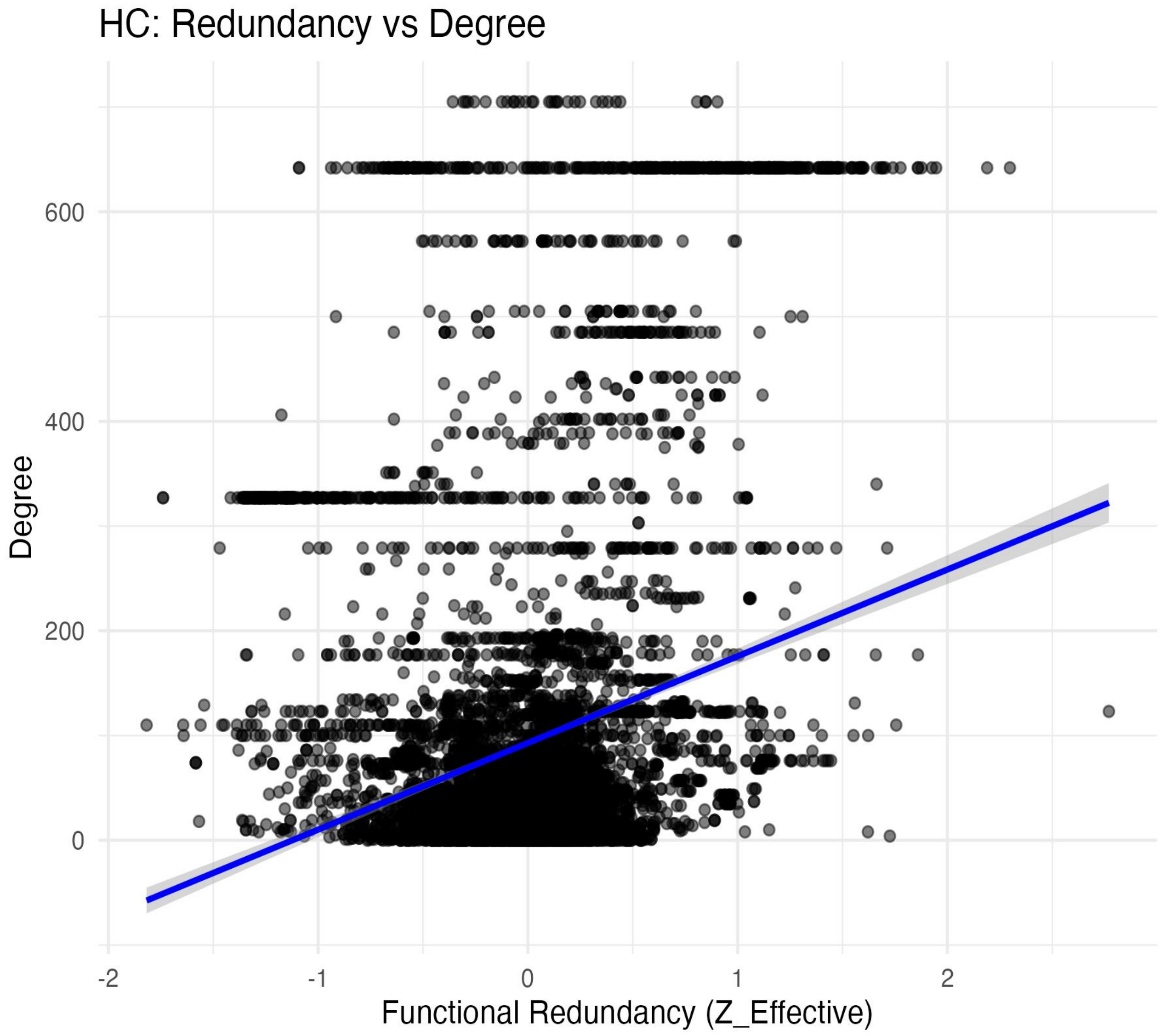


Figure S4: Healthy Control Network Analysis
Plots illustrating stable and resilient functional organization in HC microbiomes.

**Figure S5: Ulcerative Colitis Network Analysis**

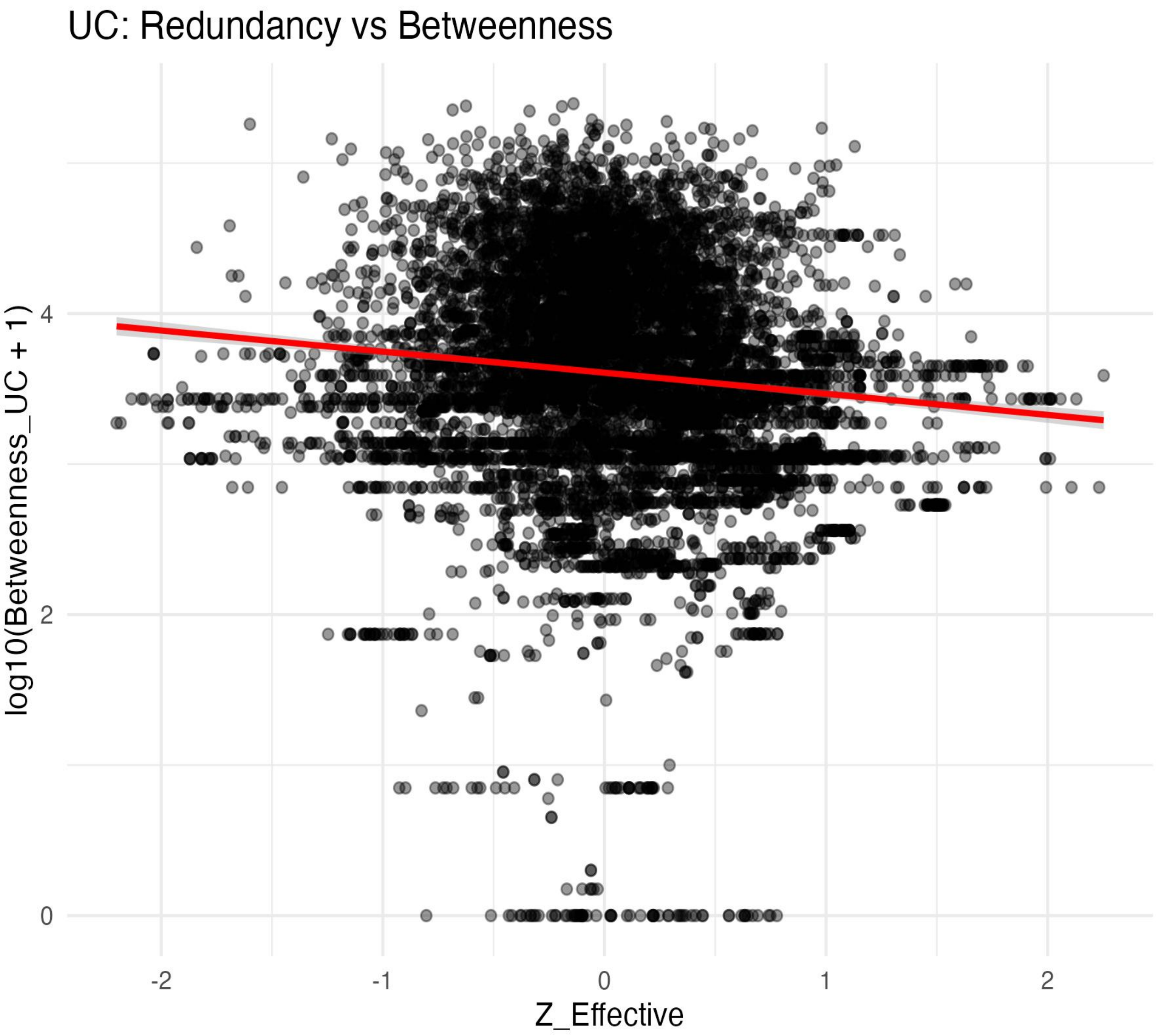

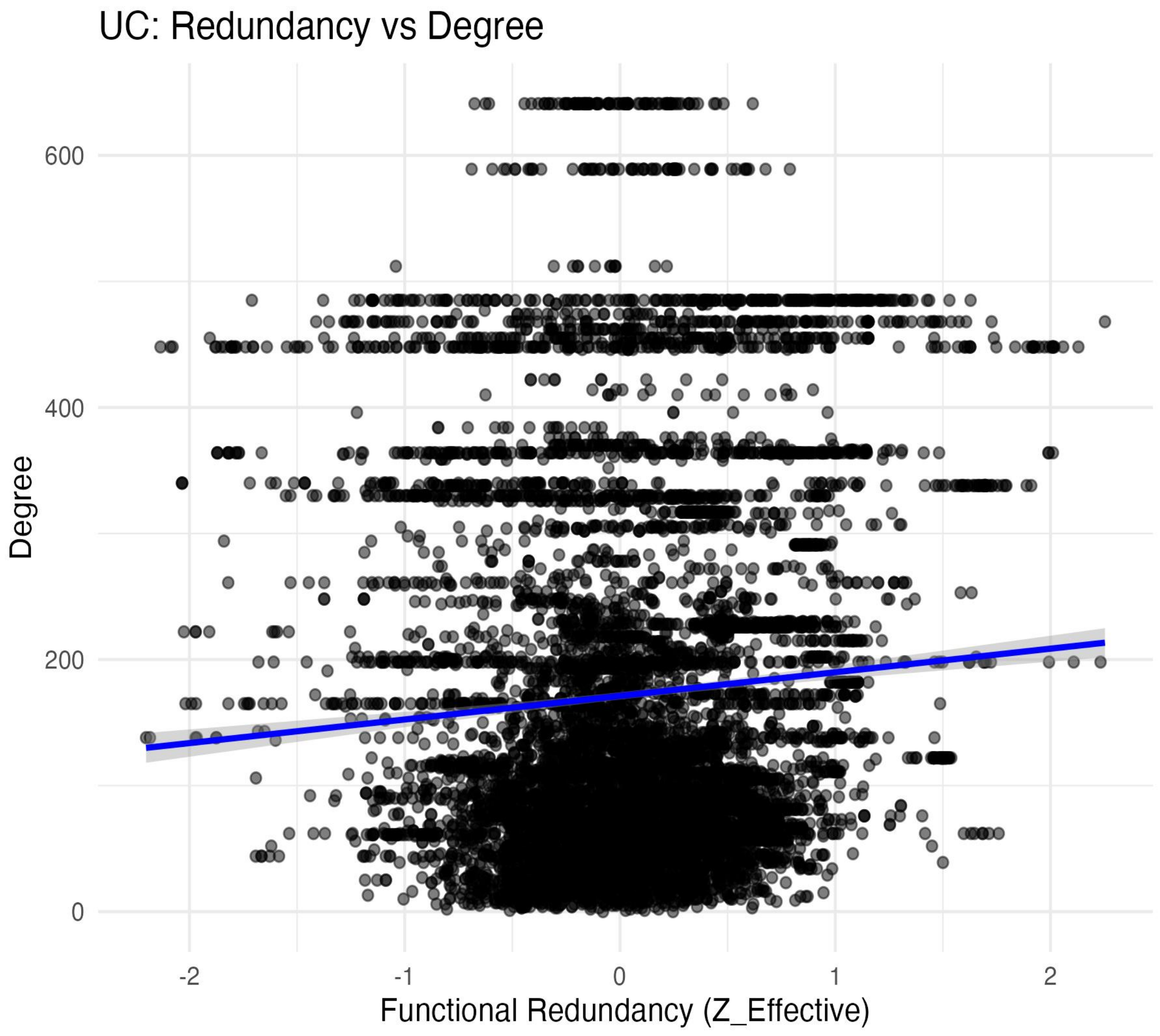


Figure S5. Network Topology in Ulcerative Colitis.
Scatter plots depicting alterations in redundancy and connectivity associated with UC.

**Figure S6: Keystone Pathways Across Disease States**

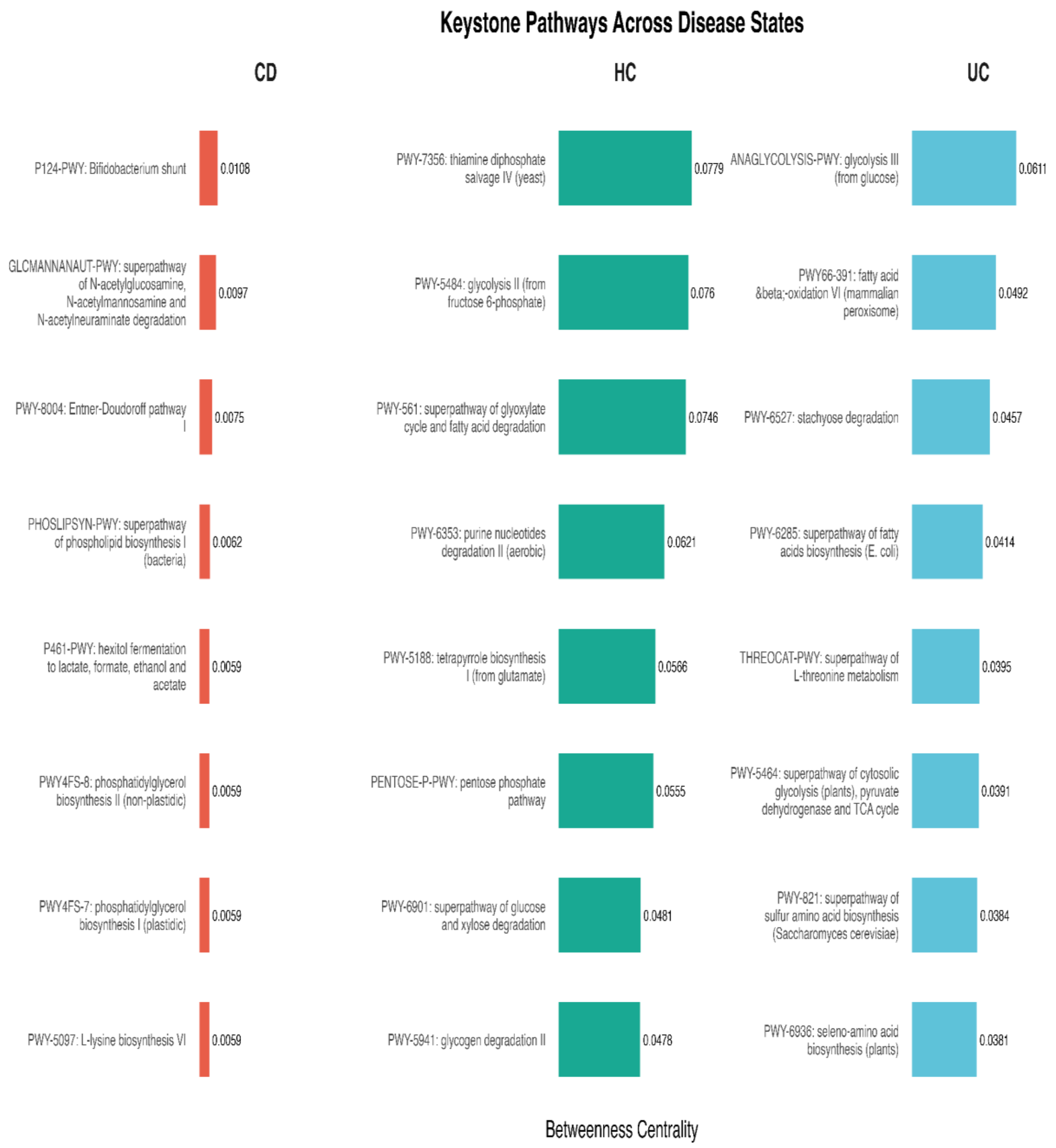


Figure S6: Keystone Pathways Across Disease States
Bar plots illustrating functional pathways with high betweenness centrality in CD, HC, and UC. These pathways represent key metabolic hubs within microbiome functional networks, highlighting disease-specific alterations in microbial functionality.

**Figure S7: Random Forest Feature Importance Analysis**

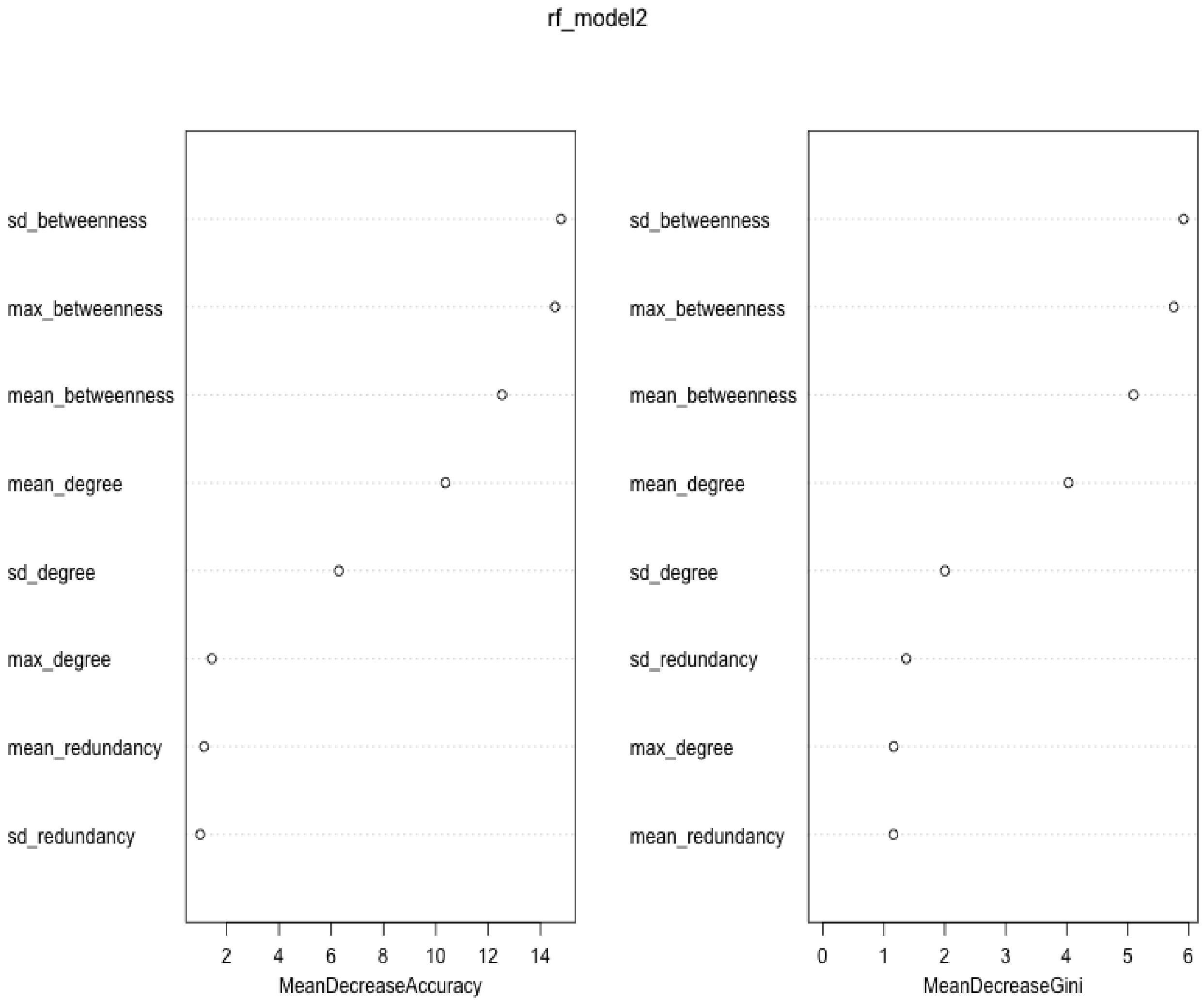


Figure S7. Random Forest Feature Importance for Disease Classification.
Variable importance plot showing the contribution of network-derived features including betweenness centrality, degree, and functional redundancy to model performance. Importance is measured using Mean Decrease Accuracy and Mean Decrease Gini, highlighting key topological predictors of inflammatory bowel disease.
