## Supplementary Materials for IBD Functional Microbiome Analysis (Datasets, Figures, and Scripts) for "NETWORK-BASED FUNCTIONAL FRAGILITY REVEALS SYSTEM-LEVEL REORGANIZATION OF THE GUT MICROBIOME IN INFLAMMATORY BOWEL DISEASE": microbiome_fragility_analysis.R

```
# ================================
# Microbiome Fragility Analysis
# ================================

# Set working directory
setwd("~/Downloads/PATHWAY_DATA-2")

# Load libraries
library(data.table)
library(tidyverse)
library(tidyr)
library(rstatix)

# ================================
# STEP 1: Read all TSV files
# ================================

files <- list.files(pattern = "*.tsv", full.names = TRUE)

read_file <- function(file) {
  df <- fread(file)
  
  colnames(df) <- c("Feature", "Abundance")
  
  df$Sample <- gsub(".tsv", "", basename(file))
  
  return(df)
}

data_list <- lapply(files, read_file)
combined_data <- bind_rows(data_list)

# ================================
# STEP 2: Clean data
# ================================

clean_data <- combined_data %>%
  filter(!grepl("^UNMAPPED", Feature)) %>%
  filter(!grepl("^UNINTEGRATED", Feature))

# ================================
# STEP 3: Keep stratified pathways
# ================================

stratified_data <- clean_data %>%
  filter(grepl("\\|", Feature))

# Split Pathway and Species
stratified_data <- stratified_data %>%
  separate(Feature, into = c("Pathway", "Species"), sep = "\\|", extra = "merge")

# ================================
# STEP 4: Normalize abundance
# ================================

stratified_norm <- stratified_data %>%
  group_by(Sample, Pathway) %>%
  mutate(Relative = Abundance / sum(Abundance)) %>%
  ungroup()

# ================================
# STEP 5: Diversity metrics
# ================================

redundancy_df <- stratified_norm %>%
  group_by(Sample, Pathway) %>%
  summarise(Redundancy = n(), .groups = "drop")

simpson_df <- stratified_norm %>%
  group_by(Sample, Pathway) %>%
  summarise(Simpson = sum(Relative^2), .groups = "drop")

shannon_df <- stratified_norm %>%
  group_by(Sample, Pathway) %>%
  summarise(
    Shannon = -sum(Relative * log(Relative + 1e-10)),
    .groups = "drop"
  )

evenness_df <- shannon_df %>%
  left_join(redundancy_df, by = c("Sample", "Pathway")) %>%
  mutate(
    Evenness = Shannon / log(Redundancy + 1e-10),
    Evenness = ifelse(Redundancy == 1, 0, Evenness)
  )

# ================================
# STEP 6: Fragility (sample level)
# ================================

fragility_df <- stratified_norm %>%
  group_by(Sample, Pathway) %>%
  summarise(
    Redundancy = n(),
    Simpson = sum(Relative^2),
    .groups = "drop"
  ) %>%
  mutate(
    Fragility_component = (1 / Redundancy) * Simpson
  ) %>%
  group_by(Sample) %>%
  summarise(
    Fragility = mean(Fragility_component),
    .groups = "drop"
  ) %>%
  mutate(
    Group = case_when(
      grepl("^HC", Sample) ~ "HC",
      grepl("^CD", Sample) ~ "CD",
      grepl("^UC", Sample) ~ "UC"
    )
  )

# ================================
# STEP 7: Pathway-level fragility
# ================================

pathway_fragility <- stratified_norm %>%
  group_by(Sample, Pathway) %>%
  summarise(
    Redundancy = n(),
    Simpson = sum(Relative^2),
    Fragility = (1 / Redundancy) * Simpson,
    .groups = "drop"
  ) %>%
  mutate(
    Group = case_when(
      grepl("^HC", Sample) ~ "HC",
      grepl("^CD", Sample) ~ "CD",
      grepl("^UC", Sample) ~ "UC"
    )
  )

# ================================
# STEP 8: Statistical testing
# ================================

pathway_stats <- pathway_fragility %>%
  group_by(Pathway) %>%
  filter(n_distinct(Group) > 1) %>%
  summarise(
    p_value = kruskal.test(Fragility ~ Group)$p.value,
    .groups = "drop"
  ) %>%
  arrange(p_value) %>%
  mutate(p_adj = p.adjust(p_value, method = "BH"))

# ================================
# STEP 9: Top pathways plot
# ================================

top_pathways <- pathway_stats %>%
  slice(1:10)

plot_data <- pathway_fragility %>%
  filter(Pathway %in% top_pathways$Pathway)

ggplot(plot_data, aes(x = Group, y = Fragility, fill = Group)) +
  geom_boxplot() +
  facet_wrap(~Pathway, scales = "free_y") +
  theme_minimal() +
  labs(title = "Top Differential Fragility Pathways")

# Save plot
ggsave("top_pathways_plot.png", width = 10, height = 6)

# ================================
# STEP 10: Effect size
# ================================

pathway_effect <- pathway_fragility %>%
  group_by(Pathway) %>%
  filter(n_distinct(Group) > 1) %>%
  kruskal_effsize(Fragility ~ Group)

# ================================
# STEP 11: Final results
# ================================

final_results <- pathway_stats %>%
  left_join(pathway_effect, by = "Pathway")

# Save results
write.csv(pathway_stats, "pathway_fragility_results.csv", row.names = FALSE)
write.csv(final_results, "final_results.csv", row.names = FALSE)

# ================================
# STEP 12: Save workspace
# ================================

save.image("my_project_workspace.RData")
```
