## Supplementary Materials for IBD Functional Microbiome Analysis (Datasets, Figures, and Scripts) for "NETWORK-BASED FUNCTIONAL FRAGILITY REVEALS SYSTEM-LEVEL REORGANIZATION OF THE GUT MICROBIOME IN INFLAMMATORY BOWEL DISEASE": README.pdf

**Project**: IBD Functional Microbiome Analysis

**Repository**: IBD_Functional_Microbiome_Supplementary

### Directory Structure:

This repository contains supplementary datasets, figures, and scripts supporting the analyses and conclusions presented in the manuscript.

**IBD_Functional_Microbiome_Supplementary/**

1. Pathway Abundance/

2. Processed Data/

2.1 Core Results/

2.2 Differential Analysis/

2.3 Shared and Unique Pathways/

2.4 Top Pathways/

3. Figures /

3.1 Supplementary_Figures.pdf

4. Scripts/

README.pdf

### 1. Pathway Abundance

This folder contains functional pathway abundance profiles generated using HUMAnN. These datasets serve as the primary input for downstream statistical, network, and machine learning analyses.

Contents

1. HC – Healthy Controls
2. CD – Crohn's Disease
3. UC – Ulcerative Colitis

File Format: .tsv

Description: Quantitative representations of microbial metabolic pathways across samples.

### 2. Processed Data

This directory includes cleaned, normalized, and analysis-ready datasets derived from pathway abundance profiles.

**Core Results**

1. final_results.csv – Consolidated dataset used for downstream analyses.
2. pathway_fragility_results.csv – Metrics describing network fragility and robustness.

**Differential Analysis**

1. CD_gain_vs_HC.csv
2. CD_loss_vs_HC.csv
3. UC_gain_vs_HC.csv
4. UC_loss_vs_HC.csv

**Shared and Unique Pathways**

1. Shared_all_pathways.csv
2. Shared_disease_pathways.csv

3. HC_only_pathways.csv
4. CD_only_pathways.csv
5. UC_only_pathways.csv

**Top Pathways**

1. FINAL_top_CD.csv
2. FINAL_top_HC.csv
3. FINAL_top_UC.csv

### 3. Figures

This folder contains all graphical outputs generated during the study. For ease of interpretation, figures are grouped thematically below. Note that these categories are descriptive and do not represent separate subdirectories.

**Network Analysis Figures**

1. Redundancy vs. Degree plots (HC, CD, UC)
2. Redundancy vs. Betweenness plots (HC, CD, UC)
3. Combined network redundancy analysis across all groups
   Example: AllGroups_Combined_Redundancy.png

**Keystone Pathway Visualizations**

1. Identification of critical pathways across disease states
   Example: Keystone_Pathways_FINAL.png

**Machine Learning Outputs**

1. Model performance comparison plots
   Example: Model_Performance_Comparison.png
2. Random Forest feature importance plots
   Example: RF_Improved_Feature_Importance.png

**Comparative Analysis Figures**

1. CD vs HC
   Example: CD_vs_HC_final.png
2. UC vs HC
   Example: UC_vs_HC_final.png
3. CD vs UC
   Example: CD_vs_UC_final_polished.png

File Formats: .png, .pdf

**Supplementary File**

1. Supplementary_Figures.pdf – Compiled supplementary figures submitted with the manuscript.

File Formats: .png, .pdf

### 4. Scripts

This directory ensures computational reproducibility of the study.

**Contents**

1. **microbiome_fragility_analysis.R** – Primary script used for data preprocessing, statistical analysis, network construction, functional redundancy analysis, and machine learning.
2. **session_info.txt** – Details of the R environment, package versions, and system configuration.

- **Machine Learning Analysis**

Machine learning models were implemented in **R** to evaluate the predictive potential of microbiome-derived functional features.

**Models Evaluated**

| Model | Algorithm | Purpose |
|---|---|---|
| Logistic Regression | Generalized Linear Model (GLM) | Baseline classifier |
| Random Forest | Ensemble Learning | Non-linear classification and feature importance |
| Support Vector Machine (SVM) | Kernel-Based Learning | High-dimensional classification |

**Model Performance**

| Model | Accuracy |
|---|---|
| Logistic Regression | 0.824 |
| Random Forest | 0.765 |
| Support Vector Machine (SVM) | 0.706 |

Logistic Regression achieved the highest predictive accuracy, demonstrating the strong discriminatory power of microbiome-derived functional features. These results are consistent with the findings reported in the manuscript.

**Software and Packages**

1. R (v4.x)
2. caret

3. ggplot2
4. dplyr
5. tidyr
6. e1071
7. randomForest

- **Software and Tools Used**

| Tool | Version | Purpose |
|---|---|---|
| Galaxy Europe | — | Bioinformatics analysis platform |
| SRA Toolkit (fasterq-dump) | v3.1.1 | Downloading sequencing data from NCBI SRA |
| FastQC | v0.11.9 | Sequencing quality assessment |
| Falco | v1.2.4 | High-performance quality control |
| fastp | v0.23.2 | Quality filtering and adapter trimming |
| Bowtie2 | v2.5.5 | Host DNA removal |
| MetaPhlAn | v4.2.4 | Taxonomic profiling |
| HUMAnN | v3.9 | Functional pathway profiling |
| R | v4.x | Statistical computing and analysis |
| ggplot2 | — | Data visualization |
| igraph | — | Network analysis |
| caret | — | Machine learning modeling |
| Random Forest | — | Classification analysis |

- **File Formats**

| Format | Description |
|---|---|
| CSV | Processed and analysis-ready datasets |
| TSV | Raw pathway abundance outputs |
| PNG | High-resolution figures |
| PDF | Supplementary figures |
| R | Analysis scripts |
| TXT | Environment and session information |
| MD | Repository documentation |

- **Reproducibility and Data Availability**

➔ Analyses were conducted using normalized pathway abundance data generated from HUMAnN.
➔ Only pathway-level functional data were used in this study.
➔ All results and figures can be reproduced using the provided scripts and datasets.
➔ The session_info.txt file ensures full computational reproducibility.

**Data Source**

All sequencing data were obtained from the NCBI Sequence Read Archive (SRA).

BioProject ID: PRJNA945504

**Preprint Information**

This supplementary repository accompanies the bioRxiv preprint:

**“**Network-Based Functional Fragility Reveals System-Level Reorganization of the Gut Microbiome in Inflammatory Bowel Disease**.”**

**Contact Information**

Mihika Kenavdekar

Bioinformatics Researcher

**License**

These supplementary materials are provided for academic and research purposes. Proper citation of the associated manuscript is required when using this data.

**Citation**

If you use these materials, please cite:

Kenavdekar, M**.,** & Natarajan, E. Network-Based Functional Fragility Reveals System-Level Reorganization of the Gut Microbiome in Inflammatory Bowel Disease. bioRxiv (Year).
DOI: To be assigned.
