## Supplementary figures and images for "NETWORK-BASED FUNCTIONAL FRAGILITY REVEALS SYSTEM-LEVEL REORGANIZATION OF THE GUT MICROBIOME IN INFLAMMATORY BOWEL DISEASE"

### AllGroups_Combined_Redundancy_vs_Betweenness.png

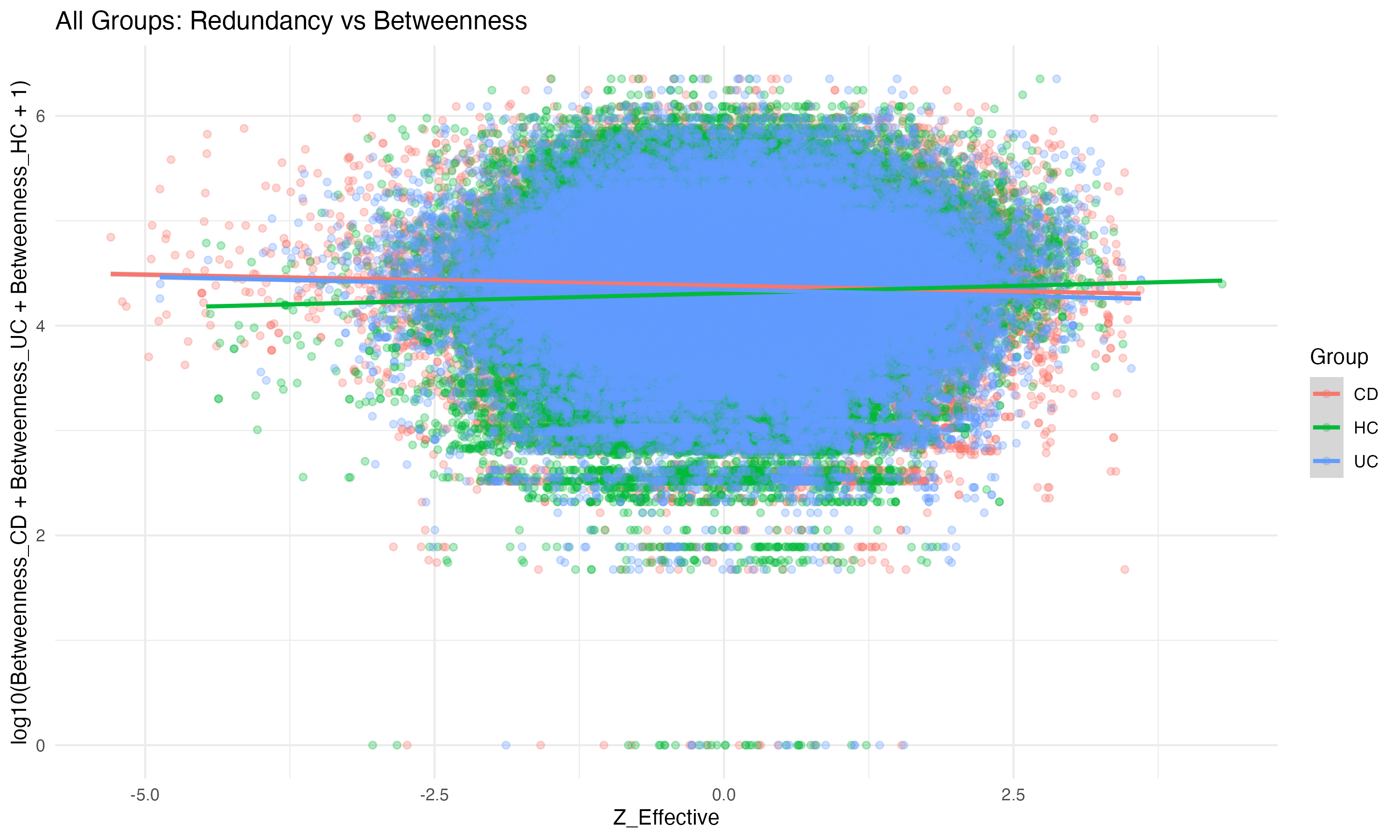

### CD_Redundancy_vs_Betweenness_log.png

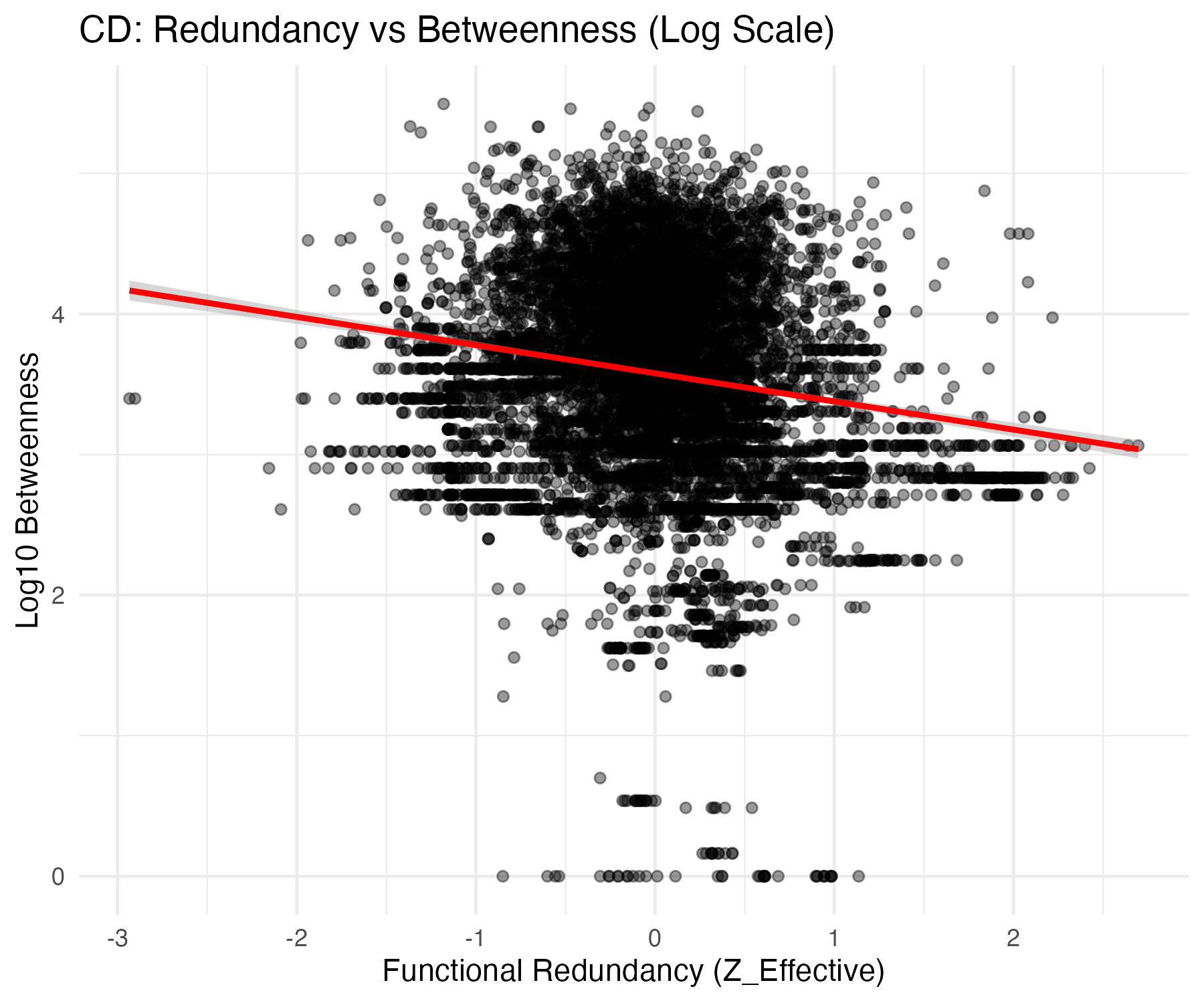

### CD_Redundancy_vs_Degree.png

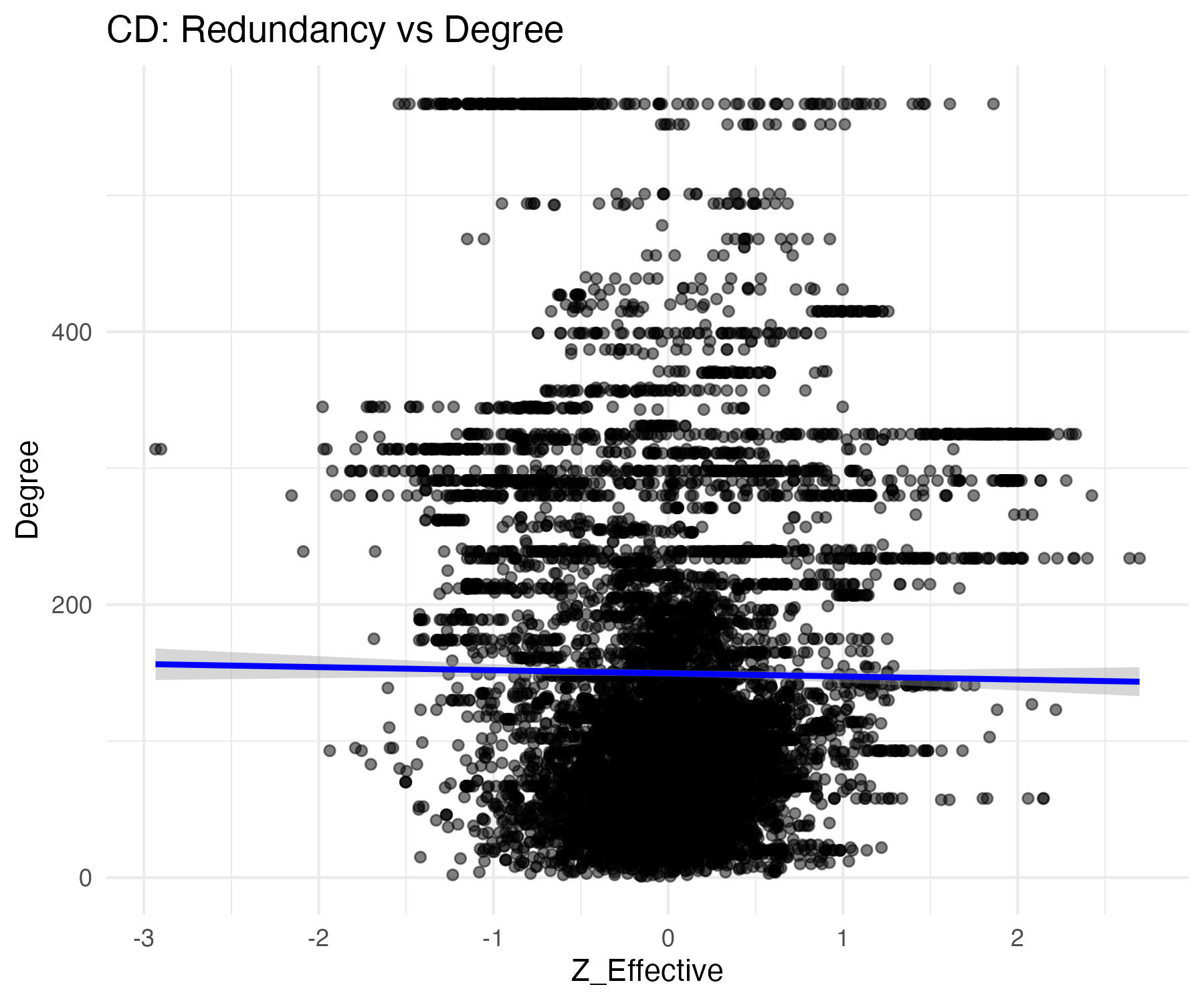

### CD_vs_HC_final.png

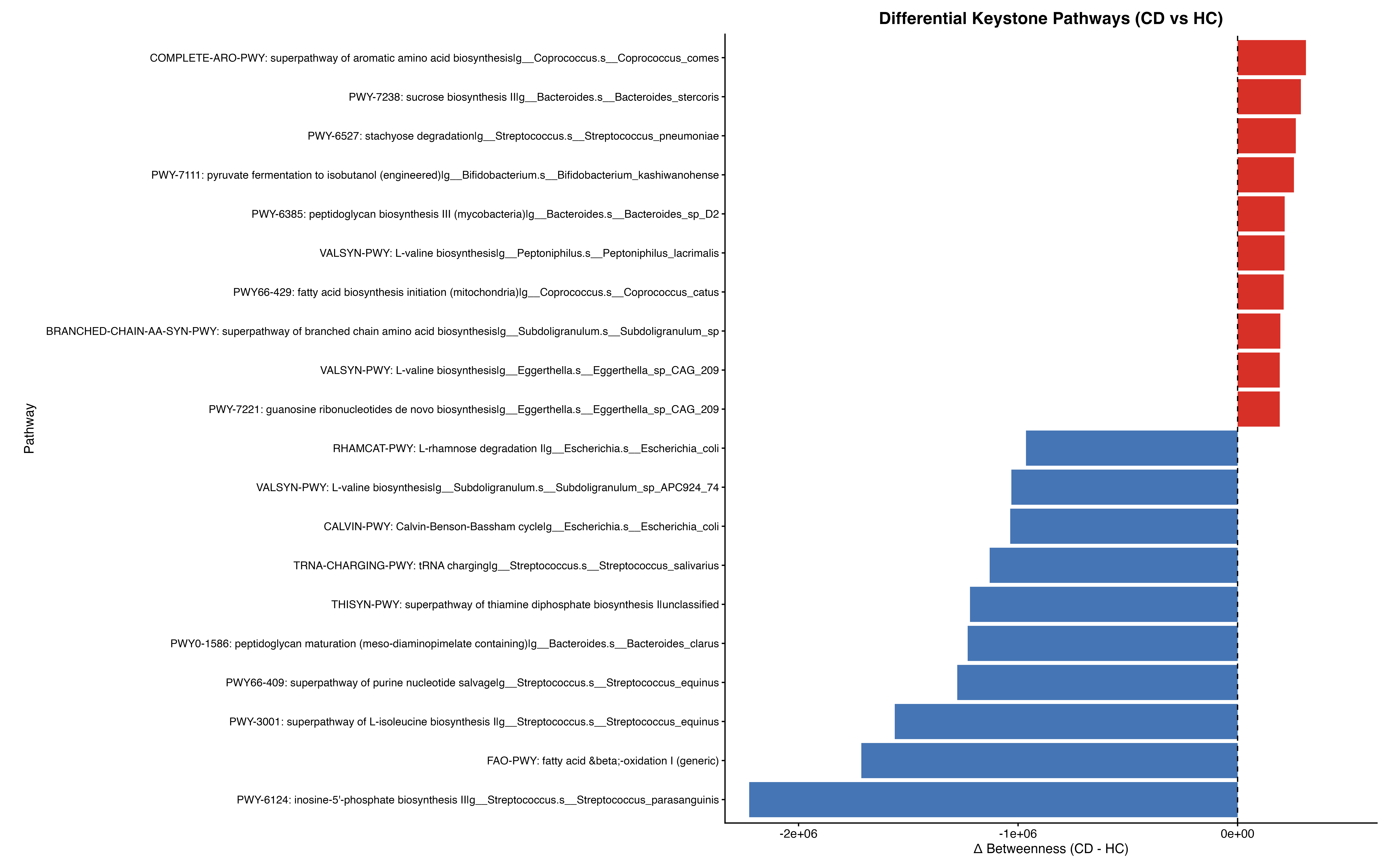

### CD_vs_UC_final_polished.png

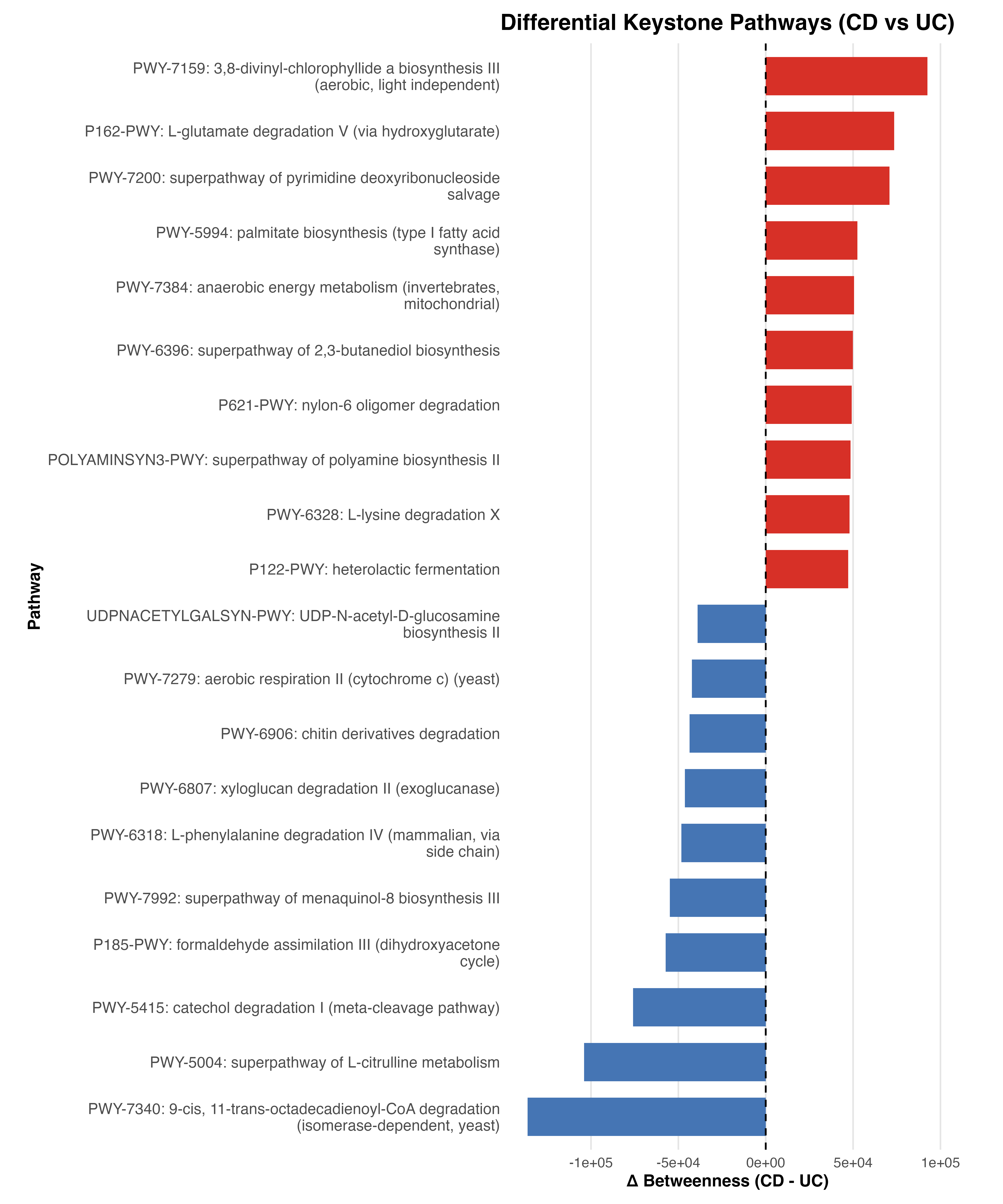

### HC_Redundancy_vs_Betweenness_log.png

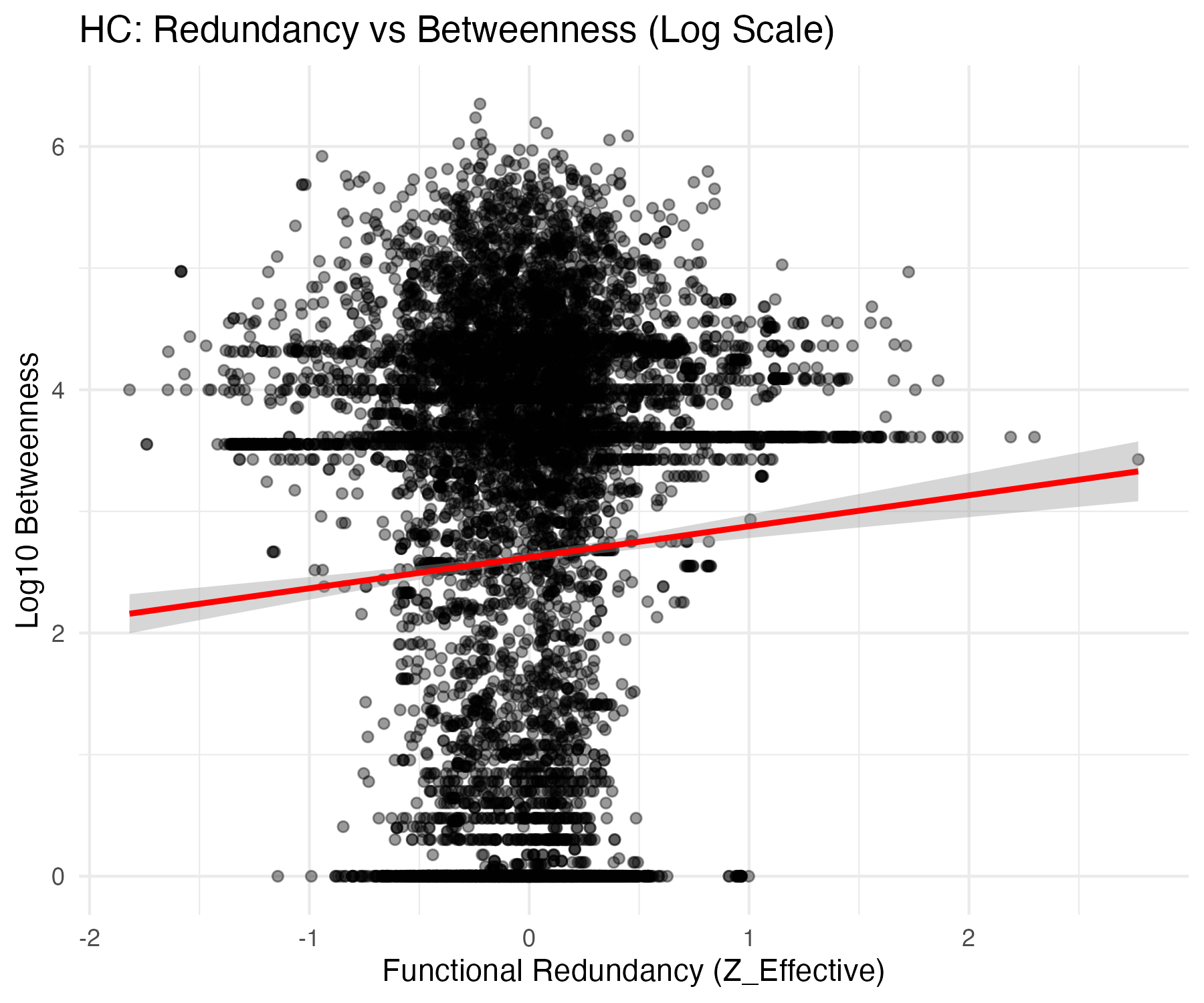

### HC_Redundancy_vs_Degree.png

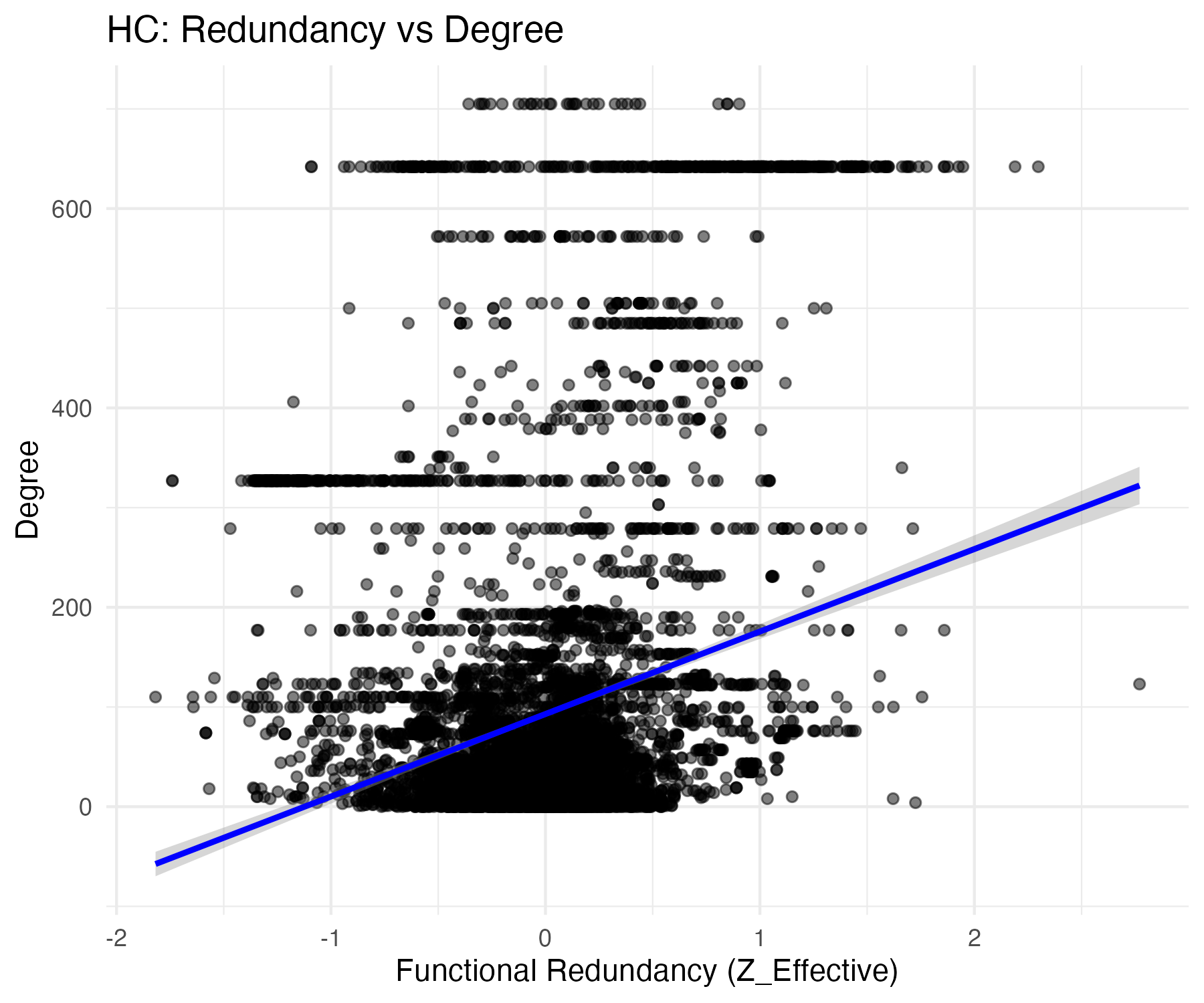

### Keystone_Pathways_FINAL_highres.png

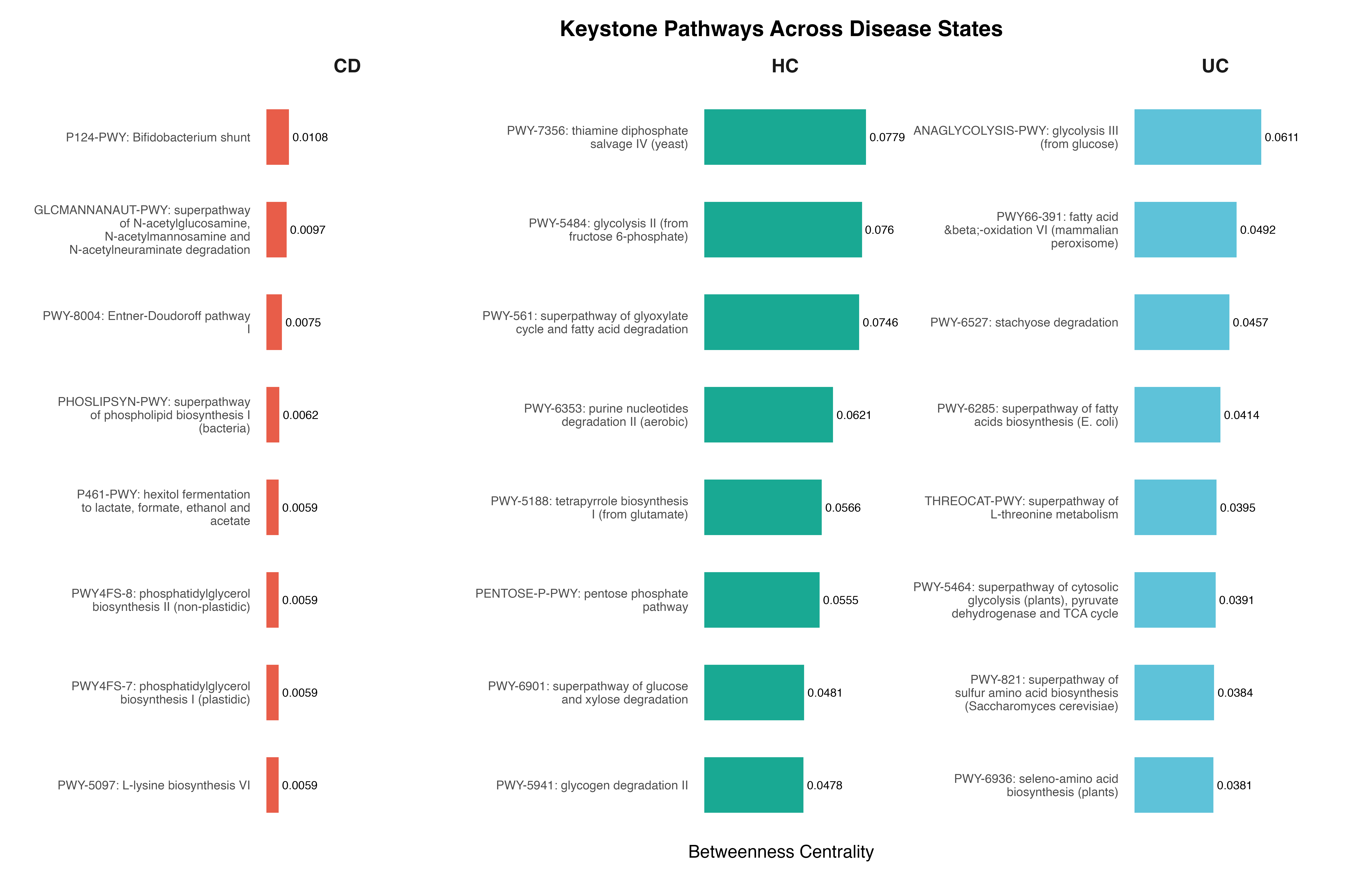

### RF_Improved_Feature_Importance.png

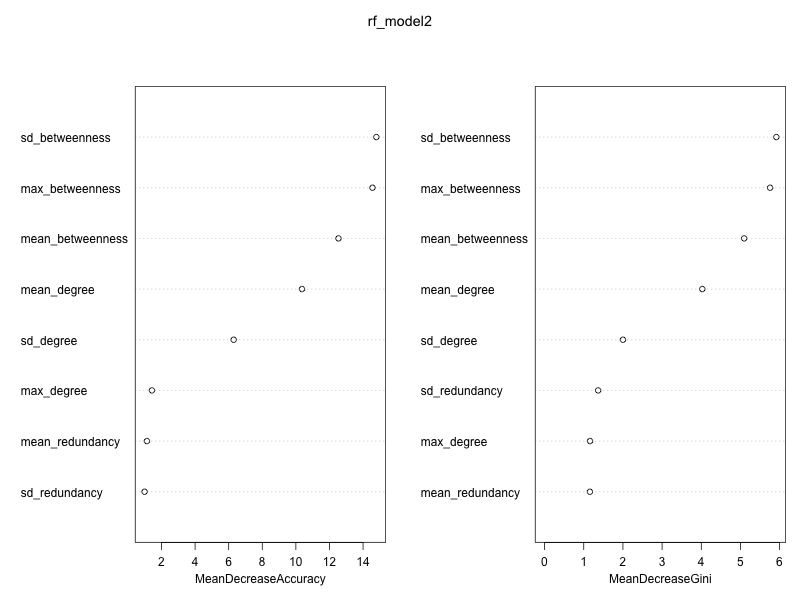

### UC_Redundancy_vs_Betweenness_log.png

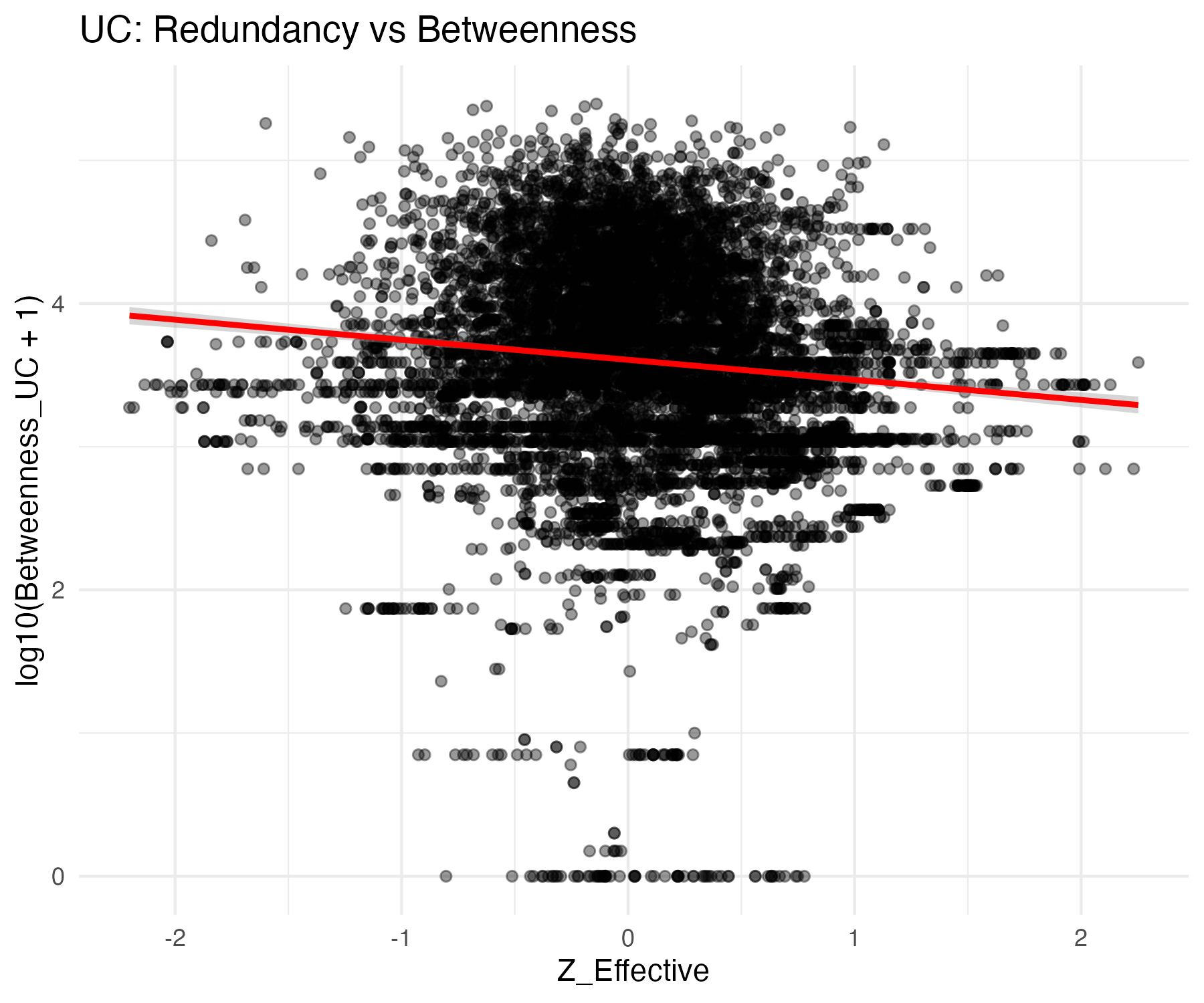

### UC_Redundancy_vs_Degree.png

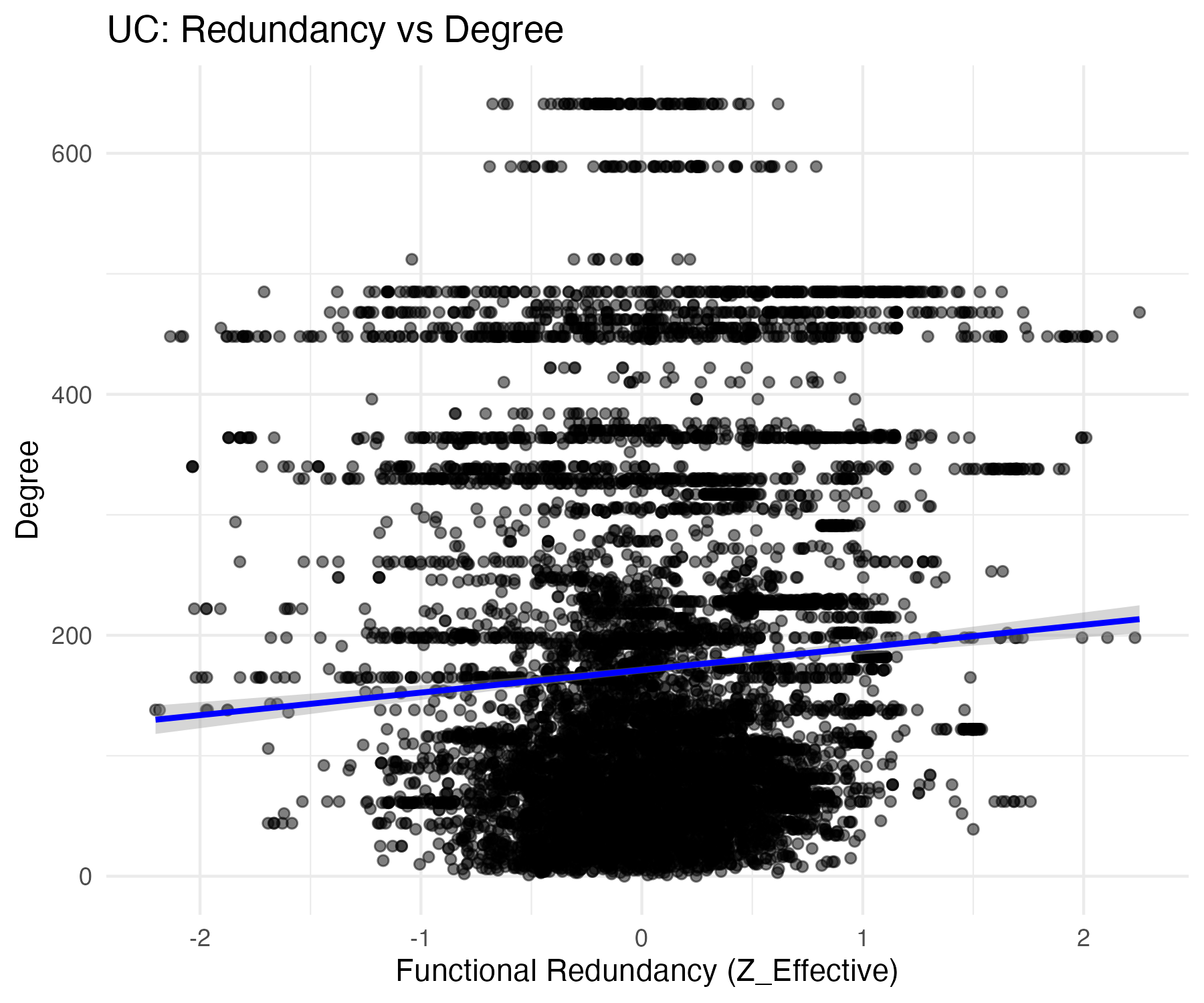

### UC_vs_HC_final.png

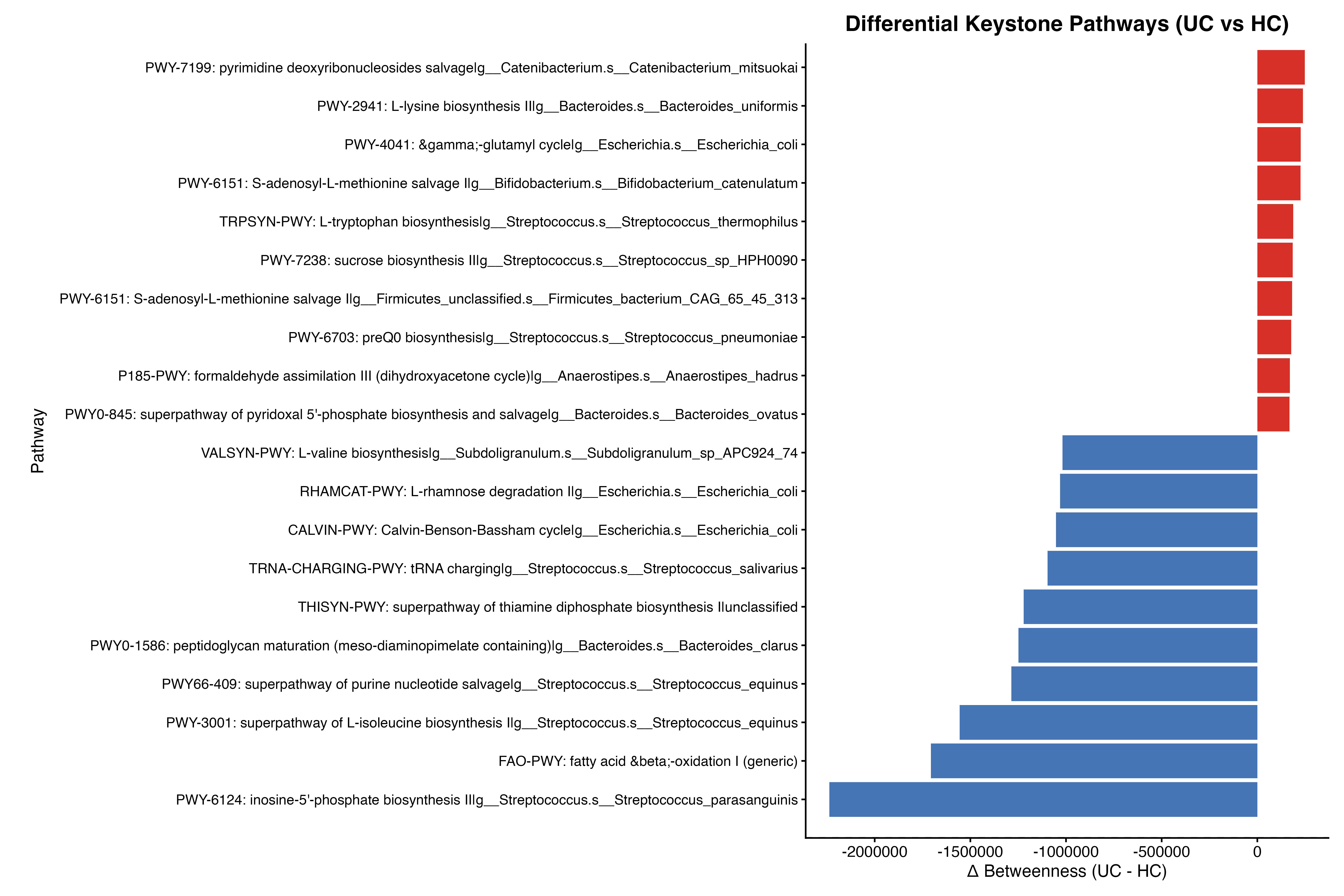
